# (2R,6R)-Hydroxynorketamine elicits rapid antidepressant effects by promoting astrocytic µ-δ opioid receptor heterodimerization

**DOI:** 10.64898/2026.05.31.729044

**Authors:** Yanxia Liang, Lingjun Wang, Yajie Li, Xiaoxue Li, Yunxiang Sun, Mengli Yang, Xuzhuo Guo, Junxu Mu, Chang Xu, Rupak Thapa, Ye Cheng, Huiqiang Zhang, Zecong He, Shi Yan, Shuo Yang, Tsz Hei Fong, Huiyuan Bai, Jia Xu, Qi Zhang, Lei Lui, Ming Li, Dongwu Xu, Wujun Geng, Jianzhong Su, Aihui Tang, Chuang Wang, Qiang Zhou, Xiang Cai

**Affiliations:** Oujiang Laboratory (Zhejiang Lab for Regenerative Medicine, Vision, and Brain Health), School of Mental Health, Wenzhou Medical University, Wenzhou, Zhejiang 325000, China; School of Basic Medical Science, Health Science Center, Ningbo University, Ningbo, Zhejiang 315211, China; School of Physical Science and Technology, Ningbo University, Ningbo 315211, China; Center of Depression, Beijing Institute of Brain Disorders, Advanced Innovation Center for Human Brain Protection, Collaborative Innovation Center for Brain Disorders, Capital Medical University, Beijing 100069, China; Anhui Province Key Laboratory of Biomedical Imaging and Intelligent Processing, HFCNS Institute of Artificial Intelligence, Division of Life Sciences and Medicine, University of Science and Technology of China, Hefei, Anhui 230026, China; College of Life Sciences, Shaanxi Normal University, Xi’an, Shaanxi 710062, China; Department of Pathology & Immunology, Baylor College of Medicine, Houston, TX, USA; State Key Laboratory of Chemical Oncogenomics, Guangdong Provincial Key Laboratory of Chemical Genomics, Peking University Shenzhen Graduate School, Shenzhen, Guangdong 518055, China; National Engineering Research Center of Ophthalmology and Optometry, Eye Hospital, Wenzhou Medical University, Wenzhou, Zhejiang 325035, China; Department of Biochemistry and Molecular Biology School of Basic Medicine, Capital Medical University, Beijing 100069, China; State Key Laboratory of Genetic Evolution & Animal Models, Yunnan Key Laboratory of Animal Models and Human Disease Mechanisms, Kunming Institute of Zoology, Chinese Academy of Sciences, Kunming, Yunnan 650201, China; School of Mental Health, Wenzhou Medical University, Wenzhou, Zhejiang 325000, China; Oujiang Laboratory (Zhejiang Lab for Regenerative Medicine, Vision, and Brain Health), Department of Anesthesiology, Wenzhou Central Hospital, Department of Pain, The First Affiliated Hospital of Wenzhou Medical University, Wenzhou, Zhejiang 325000, China

## Abstract

Ketamine produces rapid antidepressant effects but is constrained by psychotomimetic properties and abuse potential. The ketamine metabolite (2R,6R)-hydroxynorketamine (HNK) shows antidepressant-like efficacy without N-methyl-D-aspartate receptor (NMDAR) blockade, yet its upstream targets remain unclear. Here we show that HNK potentiates hippocampal excitatory transmission and reverses stress-induced behavioural deficits through opioid receptor signaling. Pharmacological and genetic analyses reveal a requirement for both µ- and δ-opioid receptors in astrocytes. Chronic stress reduces µ-δ receptor heterodimers in the hippocampus, and a single dose of HNK restores their abundance. PAINT-MINFLUX nanoscopy quantifies increased µ-δ heterodimerization, and molecular dynamics simulations indicate selective binding of HNK to the µ-receptor protomer via Asp147 and Tyr148. Mutating these residues abolishes HNK-driven heterodimer formation, downstream signaling and rapid antidepressant-like effects *in vivo*. Astrocytic µ-δ opioid receptor heterodimers thus represent a targetable mechanism for next-generation rapid-acting antidepressants.

## Main

Major depressive disorder is a leading cause of disability worldwide, and a substantial fraction of patients do not achieve remission with conventional monoaminergic antidepressants. The discovery that subanaesthetic ketamine produces rapid and sustained antidepressant effects has transformed concepts of antidepressant mechanisms and inspired the development of fast-acting treatments for treatment-resistant depression ^1–4^. However, ketamine’s broader clinical use is constrained by dissociative and psychotomimetic adverse effects and abuse liability^5,6^. Identifying upstream targets that can reproduce ketamine-like rapid efficacy without these liabilities remains a central challenge. Accumulating preclinical evidence and early-stage clinical observations indicate that (2R,6R)-HNK (hereafter referred to as HNK), a metabolite of ketamine, elicits antidepressant-like effects with minimal psychotomimetic properties^7,8^. In contrast to ketamine, HNK does not inhibit NMDARs at antidepressant concentrations^9^, suggesting that it engages an NMDAR blockade-independent pathway. Although downstream adaptations such as enhanced synaptic strength and increased expression of plasticity-related proteins have been linked to HNK^7,10,11^, the proximal molecular targets initiating its antidepressant-like effects remain poorly identified. Resolving this question is important not only for mechanistic clarity, but also for guiding the rational design of next-generation fast-acting antidepressants.

Although neurons have been the primary focus for pathophysiology of depressive illness and the antidepressant actions of HNK^7,12–14^, emerging evidence suggests that astrocyte malfunction is crucially involved in the etiology of depression ^15–17^. Beyond providing essential trophic and metabolic support for neurons, astrocytes express diverse neurotransmitter transporters and G protein-coupled receptors (GPCRs) that enable them to sense and respond to ambient synaptic activity and neuromodulators with dynamic calcium fluctuations ^18–21^. In turn, astrocytes release gliotransmitters and neurotrophic factors, such as glutamate, D-serine, ATP/adenosine, and brain-derived neurotrophic factor (BDNF) to modulate synaptic transmission and plasticity^22–25^. Such bidirectional astrocyte-neuron communication confers exquisite spatial and temporal control over synaptic activity and is essential for the maintenance of brain homeostasis^26^. Dysfunction of the synergistic cooperation between astrocytes and neurons has been implicated as a core pathological mechanism of MDD^27–29^. Nevertheless, whether HNK exerts its rapid antidepressant effects via restoration of this dysregulated signaling axis remains largely unexplored.

Among a variety of astrocytic GPCRs, opioid receptors have been increasingly implicated in emotional regulation and the response to rapid-acting antidepressants ^30–32^. Opioid receptors—including the μ (MOR), δ (DOR), κ (KOR), and nociceptin (NOPR) receptors—are expressed across brain circuits relevant to mood and can shape excitatory–inhibitory balance and neuromodulatory tone^33,34^. Beyond their canonical monomeric forms, opioid receptors can form homo- and heteromeric complexes that exhibit distinct pharmacology and signaling properties^35,36^. Among these, µ-δ opioid receptor (MOR-DOR) heterodimers are the most well-characterized and are proven to exist both *in vitro* and *in vivo*^37^. However, whether receptor heteromerization contributes to rapid antidepressant responses is largely unknown.

Here we combine hippocampal electrophysiology, behavioural assays, *in vivo* two-photon calcium imaging, cell-type-specific genetics and super-resolution single-molecule imaging to identify the upstream pathway that mediates the rapid actions of HNK. We show that HNK enhances hippocampal excitatory synaptic transmission and rapidly reverses stress-induced behavioural deficits through opioid receptor signaling that is independent of disinhibition and NMDAR inhibition. We further identify astrocytic MOR and DOR as necessary components of this response and demonstrate that stress reduces MOR-DOR heterodimers in hippocampal CA1 whereas HNK rapidly restores their abundance. Using PAINT-MINFLUX nanoscopy, we quantify HNK-driven increases in MOR-DOR heterodimerization. Through atomistic molecular dynamics simulations and targeted mutagenesis, we define key residues on the MOR protomer that are required for HNK-dependent heterodimer formation, downstream signaling and rapid antidepressant-like effects *in vivo*. Together, these findings reveal astrocytic MOR-DOR heterodimers as a mechanistic hub for the NMDAR-and disinhibition-independent rapid antidepressant actions of HNK and suggest a targetable receptor state for developing safer rapid-acting antidepressant strategies.

## Results

### HNK potentiates hippocampal excitatory transmission via opioid receptor signaling, independently of NMDAR blockade

Ketamine and HNK have been reported to enhance excitatory synaptic transmission in the hippocampus and prefrontal cortex (PFC)^7,38–40^. To elucidate the mechanism of the fast-acting antidepressant action of HNK, we compared the effect of HNK on excitatory synaptic transmission with that of ketamine in the hippocampus. We recorded excitatory postsynaptic potentials at the stratum radiatum of hippocampal CA1 by stimulation of Schaffer collateral (SC-CA1 fEPSPs) in acutely prepared hippocampal slices from naïve adult C57BL/6J mice (**Fig. 1a)**. We observed that bath application of ketamine (10 μM) or HNK (10 μM) increased the rising slope of SC-CA1 fEPSPs with similar magnitudes (62.1 ± 2.1% and 61.9 ± 5.9%, respectively) **(Extended Data Fig. 1a, b).** In addition, HNK enhanced SC-CA1 EPSCs recorded in whole-cell voltage-clamp mode **(Extended Data Fig. 1c)**. After ketamine-induced potentiation reached a steady state, subsequent bath application of HNK further elevated the slope of SC-CA1 fEPSPs by 63.1 ± 6.3% (*P* < 0.0001, ketamine+HNK versus ketamine alone) **(Fig. 1b).** In contrast, increasing the concentration of ketamine to 30 μM did not cause further potentiation of SC-CA1 fEPSPs (*P* = 0.13, 30 μM versus 10 μM ketamine) **(Fig. 1c).** MK-801 (10 μM), a selective non-competitive NMDAR antagonist, increased SC-CA1 fEPSPs slope by 95.6 ± 3.8%, and HNK perfusion further potentiated SC-CA1 fEPSPs by 53.7 ± 3.0% on top of MK-801 (*P* < 0.0001, HNK+MK-801 versus MK-801) **(Fig. 1d)**. In contrast, MK-801 mimicked and occluded ketamine-induced potentiation of SC-CA1 fEPSPs (*P* = 0.85, MK-801+ketamine versus MK-801) **(Extended Data Fig. 1d)**. These observations suggest that ketamine and HNK potentiate SC-CA1 fEPSPs via distinct mechanisms and HNK enhances excitatory synaptic transmission via an NMDAR-independent manner.

**Fig. 1.**
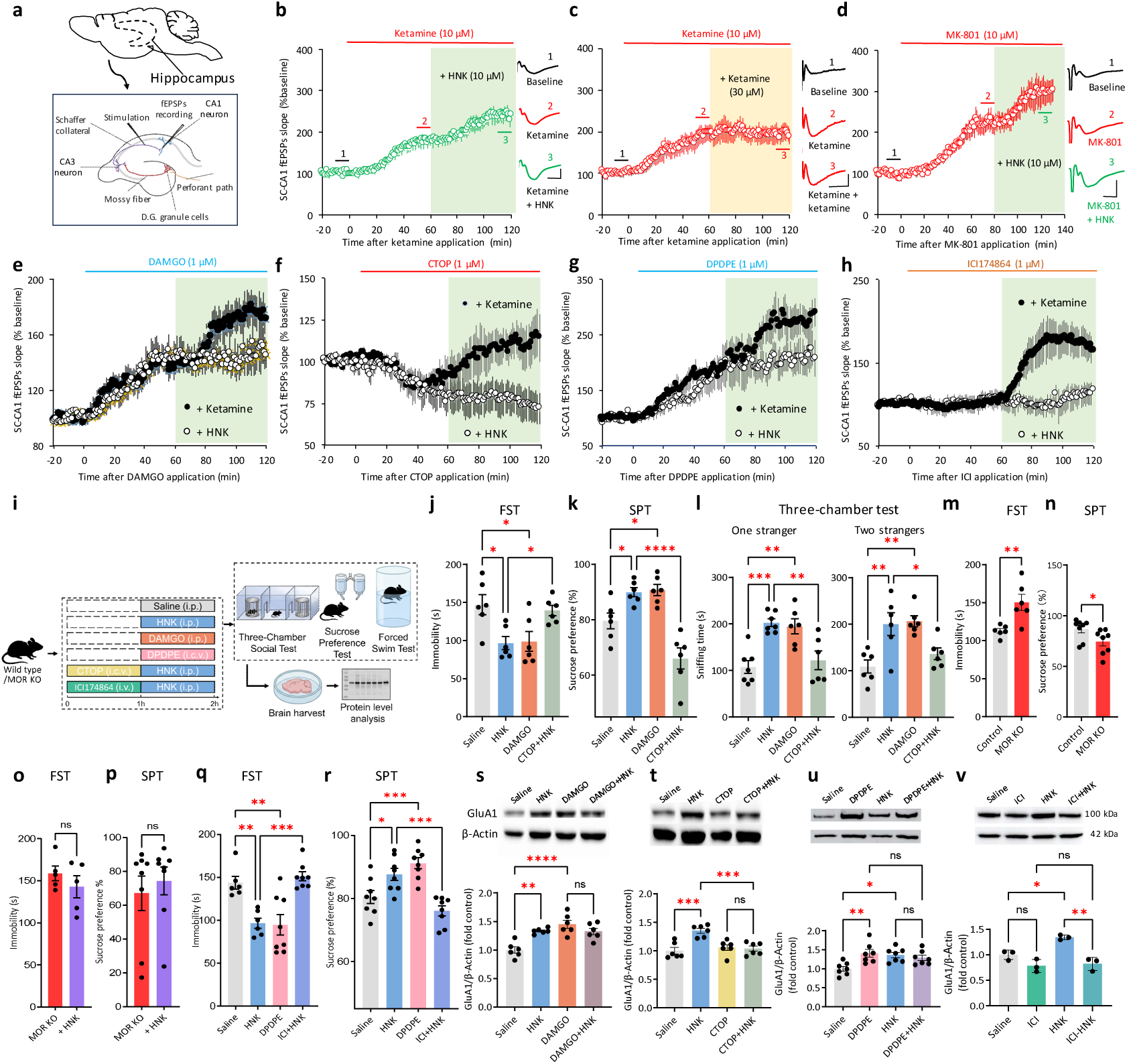
Distinct mechanisms underlie ketamine- and HNK-induced potentiation of excitatory synapses. **a,** Schematic of hippocampal slice recording: Stimulating electrode in Schaffer collaterals, recording electrode in CA1 stratum radiatum. **b,** Ketamine increased the rising slope of SC-CA1 fEPSPs elicited at CA1 stratum radiatum in acutely prepared hippocampal slices. When ketamine-induced potentiation reached a plateau, subsequent bath application of HNK further enhanced SC-CA1 fEPSPs. Left: time course of SC-CA1 fEPSPs changes; Right, representative fEPSP traces before, after ketamine treatment, and following subsequent HNK application. **c,** Increasing ketamine concentration to 30 μM did not further potentiate SC-CA1 fEPSPs after 10 μM ketamine-induced effects plateaued. **d,** MK-801, a selective and non-competitive NMDA receptor antagonist, mimicked ketamine’s potentiation of SC-CA1 fEPSPs and did not occlude the effect of HNK. **e,** DAMGO, a selective MOR agonist, increased the rising slope of SC-CA1 fEPSPs and occluded the effect of HNK on SC-CA1 fEPSPs, whereas ketamine further potentiated SC-CA1 fEPSPs on top of DAMGO. **f,** The selective MOR antagonist CTOP reduced SC-CA1 fEPSP slope and blocked the effect of HNK but not ketamine. **g**, DPDPE, a selective DOR agonist, increased the rising slope of SC-CA1 fEPSPs and occluded the effect of HNK on SC-CA1 fEPSPs, whereas ketamine further potentiated SC-CA1 fEPSPs on top of DPDPE. **h**, The selective DOR antagonist ICI blocked the effect of HNK but not ketamine. **i**, Experimental timeline for assessing the effect of MOR or DOR activation, blockade, or deletion on the antidepressant effect of HNK. **j–l,** DAMGO (10 mg·kg^-1^, i.p.) mimicked and CTOP (3 μg·kg^-1^, i.c.v.) blocked the antidepressant actions of HNK (10 mg·kg^-1^, i.p.) in FST (j), SPT (k), and three-chamber test (l). **m–p,** MOR knockout mice exhibited depression-like behaviors in FST (m) and SPT (n) and abolished the antidepressant-like effects of HNK in the two assays (FST, o; SPT, p). **q–r,** DPDPE (0.5 μg·kg^-1^, i.c.v.) mimicked and ICI (3 mg·kg^-1^, i.v.) blocked the antidepressant actions of HNK in FST (q) and SPT (r). **s–v,** Western blot analysis revealed that DAMGO and DPDPE mimicked and occluded the HNK-induced increase in GluA1 levels (s, u), whereas CTOP and ICI blocked this effect (t, v). Scale bar: 5 ms, 0.2 mV in b–d. Data are presented as mean ± SEM. \**P* < 0.05, \*\**P* < 0.01, \*\*\**P* < 0.001, \*\*\*\**P* < 0.0001. ns, not significant (statistical analyses and n values are provided in Supplementary Table 1).

Clinical and preclinical studies have shown that activation of the opioid system is required for the antidepressant effects of ketamine^41–44^, and activation of MOR has been reported to enhance excitatory synaptic transmission in the hippocampus^45,46^. To determine whether the opioid system is involved in ketamine- or HNK-induced potentiation of excitatory synaptic transmission, we tested the action of DAMGO (1 μM), a selective agonist of MOR, on SC-CA1 fEPSPs. DAMGO mimicked and occluded HNK-induced potentiation of SC-CA1 fEPSPs, whereas ketamine induced a further potentiation on top of DAMGO **(Fig. 1e, Extended Data Fig.1e).** Furthermore, CTOP (1 μM), a selective MOR antagonist, attenuated the rising slope of SC-CA1 fEPSPs and completely abolished the action of HNK, but not the of ketamine on SC-CA1 fEPSPs **(Fig. 1f, Extended Data Fig.1f)**. DOR participates in pain modulation, hedonic response and emotion control, and activation of DOR has been reported to exhibit antidepressant-like effect in animal models of depression^47,48^. DPDPE (1 μM), a selective agonist of DOR, mimicked and occluded the effect of HNK, but not that of ketamine on SC-CA1 fEPSPs (**Fig. 1g and Extended Data Fig. 1g**). The effect of HNK but not ketamine was completely abolished by ICI174864 (ICI, 1 μM), an antagonist of DOR **(Fig. 1h and Extended Data Fig. 1h).** Next, we examined the impact of opioid receptor activation or blockade on the antidepressant-like effects of HNK. In the forced swim test (FST, assessing behavioral despair), sucrose preference test (SPT, measuring anhedonia), and three-chamber interaction test (assessing sociability and social novelty), HNK rapidly (10 mg·kg^-1^, 1 h after injection) reduced immobility, increased sucrose preference, and promoted sociability and social novelty in mice, while DAMGO (10 mg·kg^-1^, i.p.) mimicked these effects and CTOP (3 μg·kg^-1^, i.c.v.) blocked these effects of HNK **(Fig. 1i–l)**. MOR knockout mice displayed depression-like behaviors in both FST and SPT, and failed to show the antidepressant-like effects of HNK **(Fig. 1m–p)**. In parallel with these observations, DPDPE (0.5 μg·kg^-1^, i.c.v.) mimicked the effects of HNK in the FST and the SPT, while ICI (3 mg·kg^-1^, i.v.) abolished these effects **(Fig. 1q–r)**. Enhancing the expression of synaptic proteins represents a crucial step for HNK-induced synaptic potentiation and synaptogenesis^7,10^. DAMGO and DPDPE mimicked the effect of HNK on the elevation of GluA1 level, whereas CTOP and ICI abolished these effects **(Fig. 1s–v)**. These observations suggest that activation of MOR- or DOR-containing opioid receptors is indispensable for HNK-induced rapid antidepressant actions.

### HNK rapidly increases the abundance of MOR-DOR heterodimers

How can the antidepressant-like actions of HNK be mimicked by activation of either MOR or DOR, and blocked by antagonists of either MOR or DOR? One possibility is that HNK facilitates the release of endogenous opioids, which subsequently activate both MOR and DOR. However, neither level of endogenous β-endorphin nor enkephalin in the hippocampus was altered by intraperitoneal injection of HNK, measured by enzyme-linked immunosorbent assay (ELISA) **(Extended Data Fig. 2a)**. Current research in the opioid field has relied heavily on the standard agonists and antagonists of opioid receptors, and the functional selectivity of these compounds is based on the assumption that opioid receptors exist as monomers or homodimers^49^. Accumulating *in vivo* and *in vitro* evidence shows that opioid receptors can form both homodimers and heterodimers^35,50,51^, and a number of classic selective MOR or DOR agonists, including morphine, DAMGO, and SNC80 activate MOR-DOR heteromeric opioid receptors with much greater efficacy than homomeric opioid receptors^49^. In light of these findings, we examined the contribution of MOR-DOR heterodimers to the antidepressant-like effects of HNK. Mice exposed to 3-week corticosterone treatment exhibited increased immobility in the FST and tail suspension test (TST), as well as reduced sociability and social novelty in the three-chamber social interaction test. A single dose of HNK (10 mg·kg^-1^, i.p.) reversed the above alterations, These effects of HNK were mimicked and occluded by pretreatment with CYM51010 (CYM, 10 mg·kg^-1^, i.p.) a known MOR-DOR heterodimer agonist) but blocked by a MOR-DOR heterodimer allosteric blocking peptide (PEP, 600 ng·kg^-1^, i.c.v.) **(Fig. 2a–d)**. CYM also mimicked the effects of HNK on the potentiation of SC-CA1 fEPSPs and elevation of GluA1 level in hippocampal slices, whereas these effects of HNK were abolished by PEP **(Fig. 2e–h).** Furthermore, Western blot analysis revealed that chronic stress decreased MOR-DOR heterodimer level and HNK restored it. The effect of HNK on MOR-DOR heterodimer was evident at 1 h, peaked at 2 h, and no significant difference was observed 24 h after injection. By contrast, acute HNK treatment did not change MOR or DOR level **(Fig. 2i–l)**. These results indicate that HNK primarily elevates the abundance of MOR-DOR heterodimers in the hippocampus, and that activation of MOR-DOR heterodimers is critical for the rapid antidepressant-like responses of HNK.

**Fig. 2.**
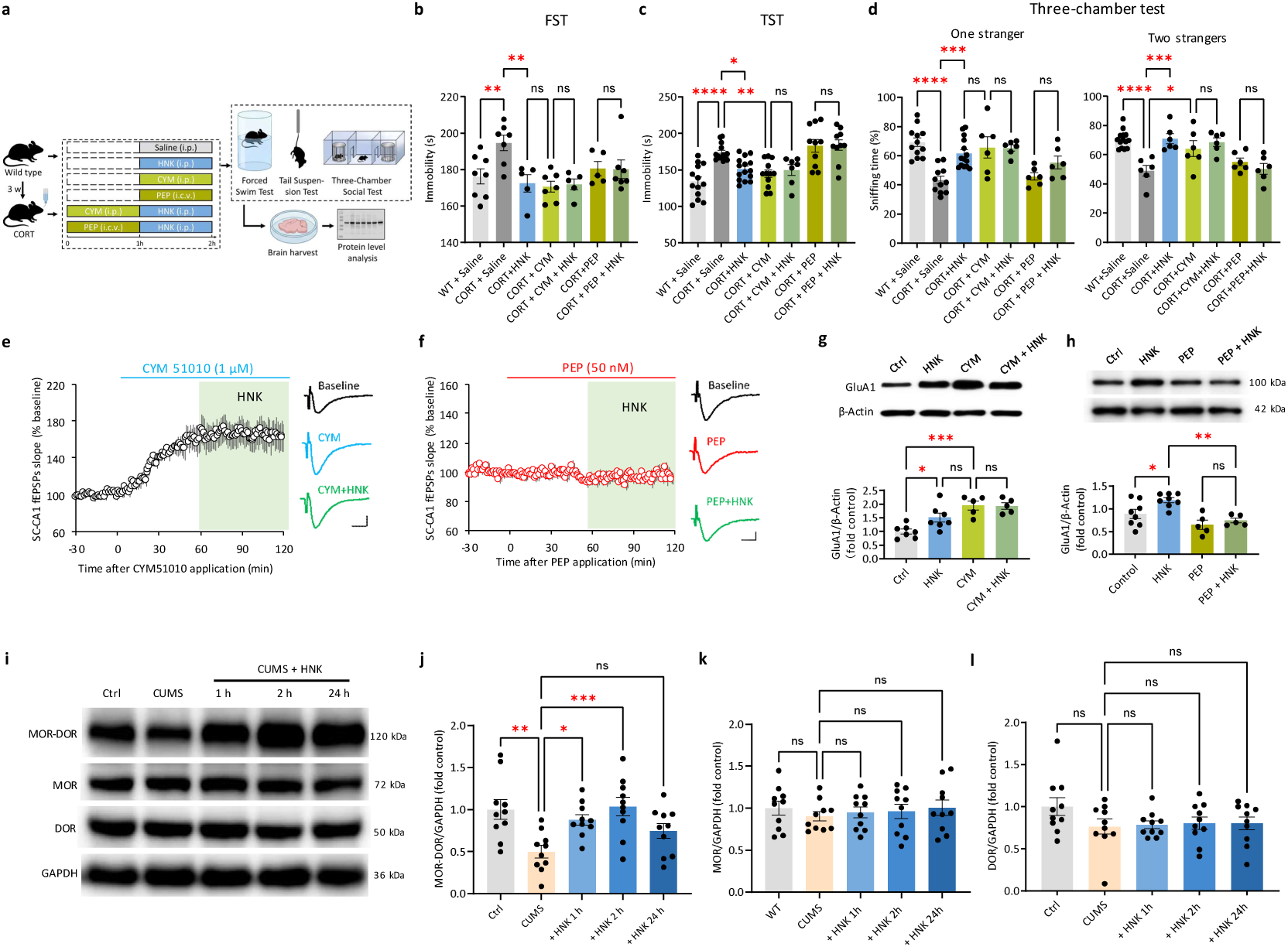
HNK increases the abundance of MOR-DOR heterodimers and blockade of the heterodimers abolishes the antidepressant-like effects of HNK. **a**, Experimental design for behavioral tests shown in b–d. **b–c**, In FST (b) and tail suspension test (c), CYM (10 mg·kg^-1^, i.p.) mimicked and occluded HNK (10 mg·kg^-1^, i.p.)-induced reduction of immobility in chronic corticosterone-treated (1 μM, in drinking water, 3 weeks) mice, and PEP (600 ng·kg^-1^, i.c.v.) blocked HNK’s actions. **d,** In three-chamber social interaction tests, CYM mimicked and occluded the antidepressant-like actions of HNK on sociability (left) and social novelty (right) tests in mice subjected to corticosterone exposure, whereas PEP abolished HNK’s effects. **e,** CYM increased the rising slope of SC-CA1 fEPSPs, and this effect mimicked and occluded HNK-induced potentiation of SC-CA1 fEPSPs. **f,** PEP (50 nM) decreased the rising slope of SC-CA1 fEPSPs and completely abolished the modulatory effect of HNK. **g,** Western blot analysis showing CYM mimicked and occluded HNK-induced elevation of GluA1 levels. **h,** PEP (50 nM) eliminated HNK-induced GluA1 upregulation. **i–l,** Using hippocampal tissue from mice subjected to different treatments, representative Western blots (i) and quantifications (j–l) show that CUMS decreased MOR-DOR level and HNK (10 mg·kg^-1^, i.p.) rapidly restored it, whereas the abundance of MOR or DOR was not significantly altered by acute HNK treatment. Scale bars: 5 ms, 0.1 mV in e, f. Data are presented as mean ± SEM. \**P* < 0.05, \*\**P* < 0.01, \*\*\**P* < 0.001, \*\*\*\**P* < 0.0001. ns, not significant (statistical analyses and n values are provided in Supplementary Table 1).

### Astrocytic MOR- or DOR-containing opioid receptors are required for the rapid antidepressant-like actions of HNK

MOR has been reported to localize at somata, dendrites, axons, and axon terminals of GABAergic inhibitory neurons including parvalbumin-expressing (PV) interneurons^52–55^, and activation of MOR on PV but not somatostatin-expressing neurons causes disinhibition of excitatory pyramidal neurons^56^. In addition, MORs are highly expressed in hippocampal astrocytes, and their activation enhances glutamate release from astrocytes, which in turn enhances excitatory synaptic transmission^46,57^. In mice, DORs are mainly expressed in GABAergic neurons and astrocytes, although weak labeling has been observed in principal neurons^58,59,60^. To determine the cellular localization of MOR-DOR heterodimers responsible for the antidepressant-like actions of HNK, we selectively ablated MOR first from PV neurons (*Oprm1*^PV^-cKO). HNK significantly reduced immobility and increased sucrose preference in both *Oprm1*^PV^-cKO and *Oprm1*^flox/flox^ control mice **(Extended Data Fig. 3a, b)**. Likewise, HNK-induced potentiation of SC-CA1 fEPSPs and increase of GluA1 level were observed in both genotypes **(Extended Data Fig. 3c, g)**. Similarly, selective deletion of DOR from PV neurons (*Oprd1*^PV^-cKO) failed to abolish the antidepressant-like actions of HNK in the FST and SPT **(Extended Data Fig. 3d, e)**, nor did it attenuate HNK-induced potentiation of SC-CA1 fEPSPs or elevation of GluA1 level **(Extended Data Fig. 3f, h)**. Consistent with these observations, the potentiating effect of HNK on SC-CA1 fEPSPs remained intact following selective blockade of GABA_A_ and GABA_B_ receptors with picrotoxin (100 μM) and CGP52432 (4 μM), respectively. **(Extended Data Fig. 3i)**. These results indicate that HNK-induced potentiation of excitatory synaptic transmission in the hippocampus is mediated neither by activation of MOR- or DOR-containing opioid receptors in PV neurons nor by disinhibition.

On the other hand, 2-h HNK incubation elevated BDNF and GluA1 abundance in hippocampal neuron-astrocyte co-cultures and increased BDNF level in pure astrocyte cultures, whereas no such upregulation was observed in neuron-only cultures **(Fig. 3a–d; Extended Data Fig. 4a–d)**. In addition, 2-h HNK treatment increased the density of GluA1-immunoreactive puncta on the proximal dendrites of neurons in the neuron-astrocyte co-cultures, but not in neuron-only cultures **(Fig. 3e–g)**. We next recorded astrocytic calcium activity in the hippocampal CA1 of head-fixed, lightly anesthetized mice via cell-specific expression of the GCaMP6f calcium indicator driven by the astrocyte-specific GfaABC_1_D promoter **(Extended Data Fig. 5a)**. Intraperitoneal injection of HNK (20 mg·kg^-1^) elicited robust calcium transients in hippocampal astrocytes **(Extended Data Fig. 5b, c)**, and these HNK-evoked astrocytic calcium responses were completely abolished by pretreatment with naloxone (1 mg·kg^-1^, i.p., 20 min prior to HNK administration) **(Extended Data Fig. 5d–f)**. To further investigate the potential modulation of neuronal activity by HNK-triggered astrocytic calcium signaling, we performed dual-color *in vivo* two-photon calcium imaging. Astrocytes were labeled with GCaMP6f, while neurons were selectively targeted with the soma-restricted red calcium indicator SomaFRCaMPi under the control of the neuron-specific human Synapsin promoter **(Extended Data Fig. 5a)**. We found that HNK induced robust calcium transients in both hippocampal astrocytes and adjacent neurons **(Extended Data Fig. 5g–I)**. Notably, HNK-elicited calcium transients peaked significantly earlier in astrocytes than in neurons **(Extended Data Fig. 5J, k)**. In addition, quantitative analysis revealed that the area under the curve (AUC) of HNK-induced calcium transients was markedly larger in neurons than in astrocytes **(Extended Data Fig. 5l)**. Intriguingly, HNK-triggered neuronal calcium elevations exhibited far longer duration than those in astrocytes. Neuronal calcium signals displayed a prominent plateau-like response followed by frequent fast transient activities, which persisted for at least 20 minutes (**Extended Data Fig. 5i, j)**.

**Fig. 3.**
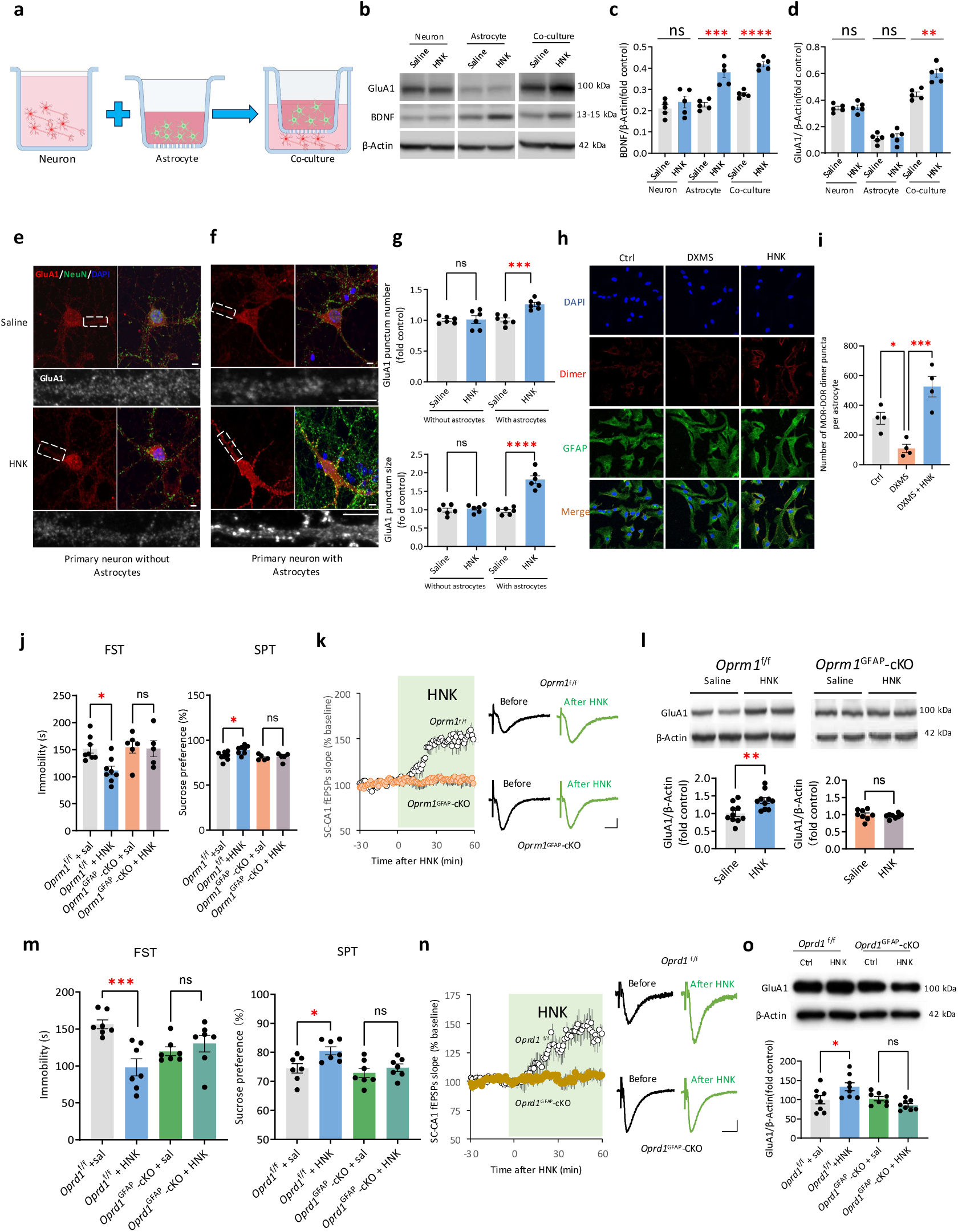
Deletion of astrocytic MOR or DOR abolishes the antidepressant-like actions of HNK. **a,** Schematic diagram of experimental procedure. **b–d,** Representative blots (b) and data summary (c, d) showing that HNK incubation (10 μM, 2 h) increased BDNF level in pure astrocyte cultures, neuron-astrocyte co-cultures, but not in pure neuron cultures (c); HNK only increased GluA1 abundance in neuron-astrocyte co-cultures (d). **e–f**, Immunofluorescence images of surface GluA1 (red) and NeuN (green) at soma and proximal processes in primary hippocampal neuron culture without (e) or with (f) astrocytes, before and after HNK incubation (10 μM, 2 h); horizontal pictures: enlargement of GluA1 clusters (boxed white). Scale bar: 2.5 μm. **g,** Quantification of number and size of GluA1 puncta in (e) and (f). **h,** Immunofluorescence images of MOR-DOR heterodimers in primary cultured hippocampal astrocytes treated with saline, dexamethasone (DXMS, 100 μM, 24 h), and DXMS plus HNK incubation (10 μM, 2 h). **i,** Quantification of number of MOR-DOR heterodimer puncta in (h). **j,** Selective deletion of MOR in astrocytes (*Oprm1*^GFAP^-cKO) blocked the antidepressant-like actions of HNK in the FST and the SPT, whereas HNK was effective in *Oprm1*^flox/flox^ controls. **k,** HNK (10 μM) enhanced SC-CA1 fEPSPs in hippocampal slices from *Oprm1*^flox/flox^ mice but not *Oprm1*^GFAP^-cKO mice. **l,** HNK increased GluA1 abundance in hippocampal slices from *Oprm1^GFAP^* controls but not *Oprm1*-cKO mice. **m,** HNK reduced immobility in the FST (left) and increased sucrose consumption in SPT (right) in *Oprd1*^flox/flox^ controls but not *Oprd1*^GFAP^-cKO mice. **n,** HNK potentiated SC-CA1 fEPSPs in hippocampal slices from *Oprd1*^flox/flox^ controls but not *Oprd1*^GFAP^-cKO mice. **o,** HNK increased GluA1 abundance in hippocampal slices from *Oprd1*^flox/flox^ controls but not *Oprd1*^GFAP^-cKO mice. Scale bars:5 μm in (e) and (f); 10 μm in (h); 5 ms, 0.1 mV in (k) and (n). Data are presented as mean ± SEM. \**P* < 0.05, \*\**P* < 0.01, \*\*\**P* < 0.001, \*\*\*\**P* < 0.0001. ns, not significant (statistical analyses and n values are provided in Supplementary Table 1).

In line with these observations optogenetic activation of hippocampal astrocytes mimicked and occluded the antidepressant-like effects of HNK in the FST and the SPT, whereas optogenetic inhibition of astrocytes abolished these effects **(Extended Data Fig. 6a, b)**. In addition, optogenetic activation of hippocampal astrocytes increased the frequency (but not amplitude) of spontaneous EPSCs (sEPSCs) recorded from CA1 pyramidal cells and occluded the effect of HNK (**Extended Data Fig. 7a–g)**, whereas optogenetic inhibition of astrocytes reduced the frequency of sEPSCs and blocked HNK’s effects (**Extended Data Fig. 7h–j).** Furthermore, HNK incubation restored dexamethasone-induced decrease of MOR-DOR heterodimers in cultured hippocampal astrocytes **(Fig. 3h–i).**

We next examined whether astrocytic opioid receptors are required for the rapid antidepressant actions of HNK. HNK failed to induce antidepressant-like effects in the FST and the SPT in mice lacking astrocytic MOR (*Oprm1*^GFAP^-cKO) **(Fig. 3j).** Similarly, HNK failed to potentiate SC-CA1 fEPSPs or elevate GluA1 level in *Oprm1*^GFAP^-cKO mice but significantly modulated these measures in *Oprm1*^flox/flox^ control mice **(Fig. 3k, l)**. In line with these findings, astrocyte-specific deletion of DOR (*Oprd1*^GFAP^-cKO) abolished the antidepressant-like effects of HNK in the FST and the SPT, whereas HNK increased sucrose preference and reduced immobility in *Oprd1*^flox/flox^ control mice **(Fig. 3m).** HNK also potentiated SC-CA1 fEPSPs and increased GluA1 level in slices from *Oprd1*^flox/flox^ mice, but not from *Oprd1*^GFAP^-cKO mice **(Fig. 3n, o).** Taken together, these results demonstrate that astrocytic MOR-DOR heterodimers are indispensable for the fast antidepressant actions of HNK.

### HNK facilitates the formation of MOR-DOR heterodimers

HNK rapidly restored the abundance of MOR-DOR heterodimers without altering total MOR or DOR levels, suggesting enhanced heterodimerization rather than upregulation of the individual subunits. To examine this possibility, we employed a bimolecular fluorescence complementation (BiFC) assay to examine the effects of HNK (10 μM) on YFP fluorescence intensity in CHO cells expressing MOR and DOR fused to the C-and N-terminal halves of YFP, respectively. HNK, but not ketamine, rapidly elevated YFP fluorescence intensity within 10 min **(Fig. 4a, b; Extended Data Movie 1)**. To directly quantify heterodimerization, we performed PAINT-MINFLUX nanoscopy^61–63^. GFP-tagged MOR and ALFA-tagged DOR were co-expressed in CHO cells and imaged sequentially via Exchange-PAINT^64^ with < 3 nm localization precision **(Fig. 4c–f)**.

**Fig. 4.**
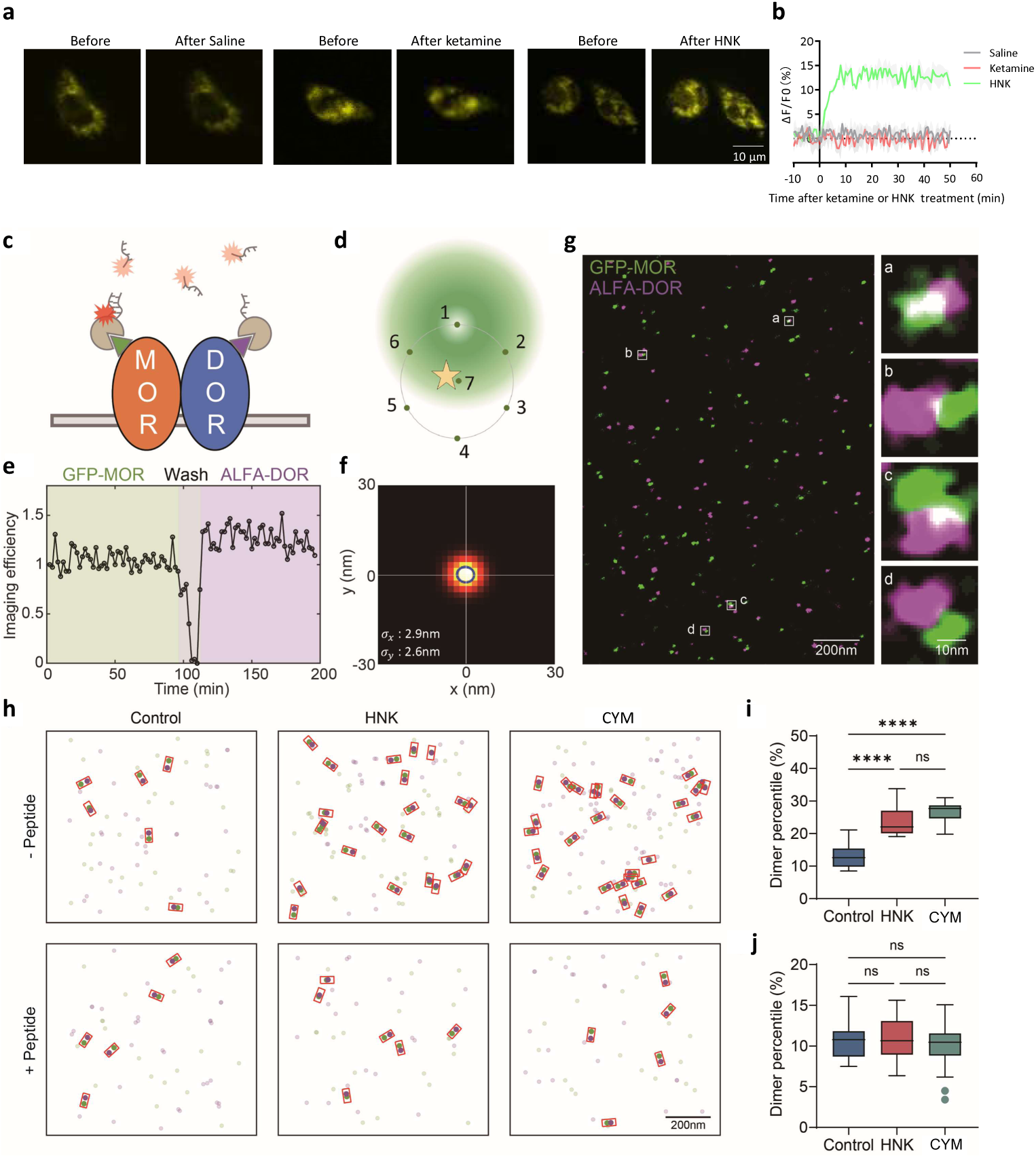
HNK promotes the formation/stability of MOR-DOR heterodimers. **a,** Representative images from bimolecular fluorescence complementation (BiFC) assays show the effects of ketamine or HNK on the fluorescence intensity of mVenus expressed in CHO cells. The coding sequences of *Oprm1* and *Oprd1* were cloned into the N-terminal and C-terminal fragments of mVenus, respectively. **b,** Time course of normalized fluorescence intensity of mVenus in the BiFC experiment shown in (a). **c,** Schematic of a putative GFP-MOR/ALFA-DOR dimer. Green and purple triangles denote the GFP and ALFA tags, respectively. **d,** Schematic of the 2D MINFLUX imaging principle. **e,** Time course of the normalized PAINT-MINFLUX single-molecule capture rate in a representative experiment. The near-zero capture rate during imager exchange indicates minimal crosstalk between the two targets. **f,** 2D histogram of localization precision for dual-target PAINT-MINFLUX. Localization precision is better than 3 nm in both x and y. **g,** Representative dual-target PAINT-MINFLUX image of GFP-MOR (green) and ALFA-DOR (purple) in CHO cells. White boxes mark putative MOR/DOR dimers; enlarged views of four examples are shown on the right. **h,** Representative MOR/DOR dimers identified in Control, HNK, and CYM groups, with and without peptide treatment. Putative dimers are outlined in red rectangles. **i–j.** Comparison of the MOR/DOR dimer fraction across the Control, HNK, and CYM groups with and without peptide treatment. Data are presented as mean ± SEM. \**P* < 0.05, \*\**P* < 0.01, \*\*\**P* < 0.001, \*\*\*\**P* < 0.0001. ns, not significant (statistical analyses and n values are provided in Supplementary Table 1).

Using a conservative 20 nm inter-receptor distance threshold for dimer calling, heterodimers were defined as paired GFP/ALFA localizations **(Fig. 4g)**. Notably, 1-h incubation with HNK or CYM significantly increased heterodimer counts, whereas PEP abolished these increases **(Fig. 4h)**. A coupled-probability dimer-calling algorithm further showed that HNK or CYM elevated the heterodimer formation probability by approximately 1.3-fold, and this effect was absent with PEP **(Fig. 4i, j)**. Collectively, these findings indicate that HNK rapidly and specifically enhances MOR-DOR heterodimerization without acutely increasing the abundance of the individual receptor subunits.

### Interrupting the direct interaction between HNK and MOR-DOR heterodimers abolishes HNK’s rapid antidepressant-like effects

We next investigated whether HNK promotes MOR-DOR heterodimerization by directly binding to the heterodimer complex. The structural stability of the MOR-DOR heterodimers, predicted by AlphaFold3 **(Extended Data Fig. 8)**, was systematically evaluated via 400 ns all-atom molecular dynamics (MD) simulations. Throughout the simulation period, the heterodimer maintained a stable transmembrane configuration, as evidenced by consistently low structural fluctuations **(Extended Data Movie 2)**. The root mean square deviation (RMSD) values for both the entire complex and each individual receptor remained below 0.5 nm, confirming well-maintained structural integrity of the heterodimer. Despite the high sequence and structural homology between MOR and DOR, HNK was observed to bind selectively to MOR within the heterodimeric complex. This binding specificity can be attributed to steric hindrance caused by the N-terminal domain of MOR, which partially occludes the orthosteric binding pocket of DOR **(Fig. 5a; Extended Data Fig. 8)**. To further assess the thermodynamic stability of the system, the time evolution of binding energy components, including polar, nonpolar, van der Waals, and total interaction energies, was analyzed using the Molecular Mechanics/Poisson–Boltzmann Surface Area (MM/PBSA) method **(Fig. 5b; Extended Data Fig. 9)**. The absence of marked fluctuations in each energy component throughout the trajectory suggests that the system had reached a reasonable thermodynamic equilibrium. Binding free energy decomposition analysis revealed that the interaction between HNK and MOR is primarily driven by a salt-bridge interaction with Asp147 (D147) and π-π stacking with Tyr148 (Y148) **(Fig. 5c, d)**. Additional residues within the binding pocket, namely Met151 (M151), Phe152 (F152), Val236 (V236), Phe237 (F237), Trp293 (W293), Ile296 (I296), His297 (H297), Val300 (V300), and Trp318 (W318), also contributed to the interaction through hydrophobic and aromatic contacts. However, their individual binding affinities were notably weaker compared to D147 and Y148, suggesting that these residues play a supportive rather than dominant role in ligand recognition.

**Fig. 5.**
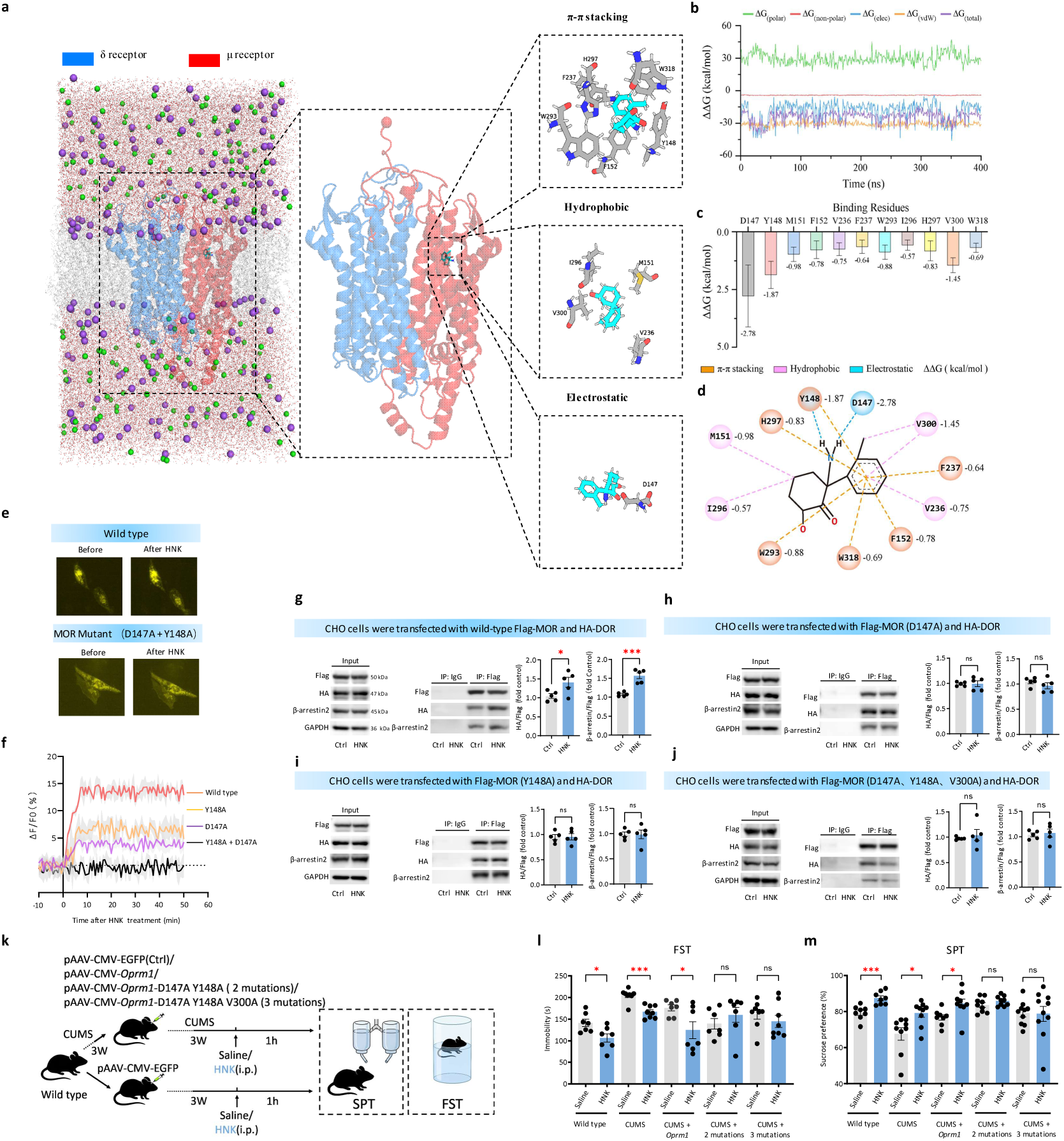
Disturbing the simulated interaction sites between HNK and MOR-DOR heterodimers abolishes the antidepressant-like effects of HNK. **a,** The binding mechanism of HNK to the MOR-DOR heterodimers was investigated through a 400 ns atomistic molecular dynamics (MD) simulation, with an explicit lipid bilayer surrounding the transmembrane region. Key interactions between HNK and the heterodimer binding pocket are categorized as π–π stacking, hydrophobic, and electrostatic interactions. **b,** Simulation equilibrium is evaluated by tracking the time evolution of binding energy, including polar, non-polar, van der Waals, and total interaction energies, using the MM/PBSA method. **c,** The residue-specific binding mechanism was further quantified by calculating the binding affinity of HNK to individual residues within the heterodimers. **d,** A 2D interaction map illustrates the binding interface between HNK and the receptor complex, highlighting key residues involved in the interaction. For all analyses, only data from the final 200 ns of the simulation are used to minimize potential bias from the initial equilibration phase. **e,** Representative images from BiFC assays demonstrate the effect of 1-h HNK (10 μM) treatment on the fluorescence intensity of mVenus expressed in CHO cells. The coding sequences of wild-type *Oprd1* (left) or mutant *Oprm1* (combined mutation of D147A and Y148A) (right) were cloned into the N-terminal and C-terminal fragments of mVenus, respectively. **f,** Time course of normalized fluorescence intensity of mVenus, where the C-terminal fragment is encoded by the wild-type *Oprd1* sequence, and the N-terminal fragment is encoded by the coding sequences of wild-type *Oprm1* or its point mutant (D147A, Y148A, or the combined mutations of D147A and Y148A). **g–j,** Co-immunoprecipitation analysis of HNK’s effect on MOR-DOR heterodimer formation and β-arrestin2 recruitment. CHO cells were transfected with Flag-tagged µ opioid receptor (Flag-MOR) and HA-tagged δ opioid receptor (HA-DOR). Left: Input lysates of CHO cells treated with HNK or vehicle (Ctrl) for 2 h were immunoblotted with antibodies against Flag, HA, β-arrestin2, or GAPDH; Middle: Lysates were immunoprecipitated with antibodies against IgG (negative control) or Flag, followed by immunoblotting with antibodies against Flag, HA, or β-arrestin2; Right: Quantification summary of the middle panel, showing the levels of HA-DOR or β-arrestin2 co-immunoprecipitated with Flag-MOR. Wild-type or point-mutated MOR sequences (tagged with Flag) were transfected into CHO cells: (g), wild-type; (j), D147A + Y148A + V300A; (h), D147A; (i), Y148A. **k,** Experimental design for behavioral tests in l–m. **l,** In the FST, HNK reduced immobility time in wild-type mice and CUMS mice infected with control AAV, and in CUMS mice infected with AAV-Oprm1, but not in CUMS mice infected with AAV-Oprm1 with 2-point mutations (D147A and Y148A), nor with 3-point mutations (D147A, Y148A, and V300A). **m,** In the SPT, HNK increased sucrose preference of wild-type mice and CUMS mice infected with control AAV, and CUMS mice infected with AAV-Oprm1, but not in CUMS mice infected with AAV-Oprm1 with 2-point mutations on D147 and Y148, nor with 3-point mutations on D147, Y148, and V300. Data are presented as mean ± SEM. \**P* < 0.05, \*\**P* < 0.01, \*\*\**P* < 0.001, \*\*\*\**P* < 0.0001. ns, not significant (statistical analyses and n values are provided in Supplementary Table 1).

To validate the accuracy of this simulation, we performed site-directed mutagenesis by substituting each predicted binding-pocket residue with alanine. In the BiFC assay, the ability of HNK to rapidly enhance the formation of MOR-DOR heterodimers was significantly blunted by MOR point mutations D147A, Y148A, or their combination (**Fig. 5e, f)**. HNK also enhanced co-immunoprecipitation of MOR and DOR as well as β-arrestin2 recruitment in CHO cells co-transfected with Flag-tagged MOR and HA-tagged DOR **(Fig. 5g)**. These effects were abolished by D147A, Y148A, or the double mutation (**Fig. 5h–j)**, whereas mutation at MOR-M151, F152, V236, W293, or H297 did not impair HNK-induced responses **(Extended Data Fig. 10)**. These results establish that D147 and Y148 serve as critical determinants for HNK binding, which in turn drives MOR-DOR heterodimerization and downstream signaling. Furthermore, overexpression of MOR carrying double (D147A/Y148A) or triple (D147A/Y148A/V300A) mutations in the hippocampus of CUMS mice completely abolished the antidepressant-like effects of HNK in the FST and the SPT **(Fig. 5k–m)**.

## Discussion

After more than two decades of intensive investigation, the mechanism underlying the antidepressant actions of ketamine remains unclear and highly controversial. A widely accepted view is that ketamine promotes the release of BDNF and enhances synaptic protein synthesis, thereby remodeling neural circuits associated with the brain’s emotion control system that are disrupted in depression^2,4,65–67^. Current debates on the antidepressant mechanism of ketamine primarily focus on whether the antidepressant effects of ketamine are initiated by NMDAR blockade or other non-NMDAR-mediated signaling such as that mediated by its metabolites including HNK^2,4,7,29,68,69^. In the current study, we demonstrate a fundamental distinction in the mechanisms engaged by ketamine and HNK in potentiating excitatory synaptic transmission in the hippocampus. This divergence indicates that ketamine and its metabolites may work collectively to restore malfunctioning excitatory synapses in depression. A key finding of our study is the critical role of astrocytic MOR-DOR heterodimers in mediating HNK’s rapid antidepressant effects. By demonstrating that HNK enhances the formation/stabilization of these heterodimers through a direct interaction with the MOR subunit, and that this interaction is necessary for HNK’s behavioral and synaptic effects.

Several aspects of our study extend previous research on the antidepressant mechanisms of ketamine and its metabolites. First, our work identifies a specific subpopulation of opioid receptors—astrocytic MOR-DOR heterodimers—as the key mediator of HNK’s actions. This specificity is important, as it provides a potential target for the development of new antidepressants that can mimic HNK’s rapid effects without the side effects associated with non-selective opioid receptor activation. Second, the application of advanced imaging techniques such as PAINT-MINFLUX nanoscopy allowed us to directly visualize and quantify the effect of HNK on MOR-DOR heterodimer formation with high spatial precision, providing direct evidence for HNK’s role in promoting heterodimerization. Third, we demonstrated that HNK-induced potentiation of excitatory synaptic transmission is independent of disinhibition, since blockade of GABA receptors failed to abolish the effect of HNK, which possesses an essential distinction from a recent finding regarding the action of ketamine on MOR (μ-opioid receptor) in SST neurons to exert its antidepressant effect^13^. Finally, our MD simulations and site-directed mutagenesis studies provide a detailed molecular understanding of HNK’s interaction with the MOR-DOR heterodimer, which could guide the rational design of novel compounds that target this heterodimer with high affinity and selectivity.

Accumulating evidence has demonstrated that maladaptive astroglia-neuron interaction is involved in the aetiology of depression^18,29,70,71^. Our *in vivo* two-photon calcium imaging experiments reveal that HNK evokes robust naloxone-sensitive calcium responses in hippocampal CA1 astrocytes. Importantly, HNK-evoked calcium transients peak significantly earlier in astrocytes than in neurons, suggesting that astrocytes may serve as the primary cellular target of HNK. Astrocyte-neuron co-culture experiments and optogenetic experiments suggest that HNK acts on astrocytic MOR-DOR heterodimers to trigger downstream signaling pathways that promote the release of gliotransmitter or astrocyte-derived neurotrophic factors (such as BDNF), and subsequently enhance the expression of synaptic proteins in neurons, ultimately leading to the restoration of excitatory synaptic function and the alleviation of depressive-like behaviors. Astrocyte-specific MOR or DOR knockout mice, combined with fluorescent probes targeting glia-derived transmitters and neuromodulators, will facilitate future investigation into the detailed mechanisms by which HNK rescues the impaired astrocyte–neuron bidirectional communication under depressive conditions.

Together, our findings demonstrate that HNK restores excitatory synaptic transmission and plasticity by directly promoting astrocytic MOR-DOR heterodimerization at the astrocyte-neuron interface in the hippocampus. These results not only deepen our understanding of the pathophysiology of depression but also highlight a promising target for the development of next-generation, fast-acting antidepressants with improved efficacy and safety profiles.

## Materials and Methods

### Animals

C57BL/6J mice were purchased from Beijing Vital River Laboratory Animal Technology Co., Ltd., Beijing, China. MOR KO mice (007559), *Oprm1*^flox/flox^ mice (030074), and PV-Cre mice (017320) were purchased from the Jackson Laboratory, Bar Harbor, ME, USA. *Oprd1*^flox/flox^ mice (#NM-CKO-220401) and GFAP-Cre mice (#NM-KI-242462) were purchased from Shanghai Model Organisms Center, Inc., Shanghai, China. *Oprm1*^PV^-cKO and *Oprm1*^GFAP^-cKO mice were generated by crossing *Oprm1*^flox/flox^ mice with PV-Cre mice or GFAP-Cre mice. *Oprd1*^PV^-cKO and *Oprd1*^GFAP^-cKO mice were generated by crossing *Oprd1*^flox/flox^ mice with PV-Cre mice or GFAP-Cre mice. Mice were housed in ventilated cages (5-6 mice per cage) under standard laboratory conditions (23 ± 1°C, humidity of 50%-60%, a 12/12 h light/dark circadian cycle with access to food and water ad libitum). All animals were habituated for a week to adapt to the housing conditions after purchase and then subjected to different training or drug treatments. Mice were randomly numbered using a table of numbers generated by Microsoft Excel and assigned to different treatment groups. Both male and female mice used for tissue preparation or behavioral tests in this study were 8 to 14 weeks old. All experimental procedures involving animals were performed in accordance with the Biological Research Ethics Committee of Oujiang Laboratory, Wenzhou, Zhejiang, China (Permit No.: AEEI-2019-133).

### Chemicals

(R,S)-ketamine provided by Dr. Sungchil Yang of City University of Hong Kong, (2R,6R)-HNK (Tocris), DAMGO (Tocris), CTOP (Tocris), MK-801 (MCE), D-APV (MCE), DPDPE (MCE), ICI174864 (MCE), Picrotoxin (MCE), CYM51010 (MCE), CGP52432 (Millipore), and MOR-DOR heterodimer allosteric inhibitor peptide (PEP; amino acid sequence: VTACTPSDGPGGGAAAYGRKKRRQRRR) (QYAOBIO, Shanghai, China) were dissolved in 0.9% saline to make a stock solution. Corticosterone (Biosharp, Hefei, Anhui, China) was dissolved in 100% ethanol to prepare a stock solution. In behavioral tests, (R,S)-ketamine (10 mg·kg^-1^), HNK (10 mg·kg^-1^), DAMGO (10 mg·kg^-1^), and CYM51010 (10 mg·kg^-1^) were administered intraperitoneally (i.p.), while CTOP (3 μg·kg^-1^), DPDPE (0.5 μg·kg^-1^), and PEP (600 ng·kg^-1^) were administered intracerebroventricularly (i.c.v.). ICI174864 (3 mg·kg^-1^) were administered intravenously (i.v.). Corticosterone was added to the drinking water of test mice. For electrophysiology, immunoblotting, and immunoprecipitation experiments using brain slices, drugs were diluted in ACSF to a final concentration as indicated in the results or figure legends (please see extended data table 3 for the catalog numbers of key chemicals).

### Hippocampal slice preparation

This method has been described in detail in our previous studies^6,38,72,73^. Briefly, mice were decapitated with a guillotine after being anesthetized with isoflurane (RWD Life Science, Shenzhen, Guangdong, China). Mouse brains were rapidly removed, and hippocampal dissection was performed in ice-cold artificial cerebrospinal fluid (ACSF: 120 mM NaCl, 3 mM KCl, 1 mM NaH_2_PO_4_, 1 mM MgCl_2_, 2 mM CaCl_2_, 25 mM NaHCO_3_ and 20 mM glucose) bubbled with 95% O_2_/5% CO_2_. Subsequently, isolated hippocampi were sectioned coronally into slices of 400 μm thickness using a vibratome (#VT1200S, Leica Microsystems, Wetzlar, Hesse, Germany) and kept in a submerged incubation chamber with oxygenated ACSF at 35 ± 0.5°C for 30 min, then incubated continuously at room temperature (23 ± 1°C) for 1 h before recording.

### Extracellular field potential recording

After incubation, hippocampal slices were transferred to a submersion-type recording chamber and perfused with ACSF (flow rate: 1-2 mL·min^-1^) at room temperature. Concentric bipolar tungsten electrodes were placed in the stratum radiatum subfield of CA1 to stimulate Schaffer collateral afferents. Extracellular recording pipettes (3-5 MΩ, World Precision Instruments, Sarasota, FL, USA) were filled with ACSF and placed 200-350 μm from the stimulating electrodes in stratum radiatum. Stimuli (100 ms duration) were delivered at 0.05 Hz. The stimulus intensity was set at 150% threshold intensity, resulting in field EPSPs (fEPSPs) of 0.1-0.2 mV. All compounds were applied by perfusion. Field EPSPs were recorded using MultiClamp 700B amplifier (Molecular Devices, San Jose, CA, USA). Signals were amplified 100× and filtered at 10 kHz before digitization with a Digidata 1550B A/D converter (Molecular Devices). Data were collected and analyzed with Clampex 11.3 software (Molecular Devices). In some experiments, Picrotoxin (100 μM) and CGP (4 μM) were included in ACSF to block GABA_A_ and GABA_B_ receptors, respectively.

### Whole-cell patch-clamp recording

Whole-cell voltage-clamp recordings were obtained with patch pipettes (World Precision Instruments) filled with 135 mM CsCH_3_SO_3_, 10 mM HEPES, 10 mM NaCl, 1 mM MgCl_2_, 0.1 mM BAPTA, 2 mM Mg^2+^-ATP, 10 mM phosphocreatine, and 5 mM QX-314 (#1014/100, Tocris Bioscience), with pH adjusted to 7.25-7.30 by CsOH, and with final osmolarity adjusted to 295-300 mOsm·L^-1^. Hippocampal CA1 pyramidal cells were visualized with a 40× water-immersion objective and infrared optics on an upright microscope (#BX51WI; Olympus Corporation, Tokyo, Japan).

Images were detected with an IR-sensitive charge-coupled device (CCD) camera (ORCA-Fusion, Hamamatsu Photonics K.K., Hamamatsu, Shizuoka, Japan). Electrode resistances in the bath were 4-7 MΩ, and series resistances of < 40 MΩ were accepted. Data were collected using a Multiclamp 700B amplifier (Molecular Devices), low-pass-filtered at 2 kHz, and digitized at 5 kHz using a Digidata 1550B A/D converter and Clampex 11.3 software (Molecular Devices). The series resistance and capacitance were compensated automatically after a stable gigaseal was formed.

### Chronic unpredictable mild stress (CUMS)

Mice were randomly divided into a control group and a CUMS group. Animals in the CUMS group were treated as follows: Day 1: strobe light (1 h), restraint (30 min), and food deprivation (16 h); Day 2: social isolation (16 h) and cage tilting (45°, overnight); Day 3: food and water deprivation (16 h), strobe light (1 h), and wet bedding (overnight); Day 4: restraint (30 min), strobe light (1 h), and overnight illumination; Day 5: cold water swim (8–10°C, 3–5 min), strobe light (1 h), and water deprivation (16 h); Day 6: tail clipping (at the base, 10 min), cage tilting (45°, overnight), and wet bedding (overnight); Day 7: restraint (30 min) and water deprivation (16 h). This cycle was repeated for 3–5 weeks. The control group was not disturbed except for necessary routine procedures.

### Forced swim test (FST)

The FST was performed as described in our previous study^38^ with minor modifications. Mice first underwent a 15-min pretest session, during which they were individually placed in a vertical Plexiglas cylinder (4000 mL total volume) filled with 3000 mL of water maintained at 20-23°C. Water was refreshed between subjects to ensure consistency. Upon completion of the pretest, mice were gently dried with a towel and returned to their home cages. The test session was conducted 24 h after the pretest, lasting 6 min. Immobility duration—defined as the total time spent motionless or floating, excluding minimal movements required to maintain balance—was quantified during the final 4 min of the test using automated analysis software (ANY-maze 7.3, Global Biotech, Inc., USA). A reduction in immobility time (i.e., increased swimming or struggling behavior) was interpreted as an antidepressant-like response.

### Sucrose preference test (SPT)

The SPT was performed according to our previous publication^38^ with minor modifications. Mice were habituated to two water bottles for the first two days, with bottle positions switched every 24 h. Over the next two days, one bottle contained water and the other 1% sucrose solution. On day five, mice were water-deprived for 6 h, and the initial weights of both bottles were measured. They were then exposed to water and 1% sucrose in the dark for 24 h, with bottle positions switched at the 12-h mark. Final consumption was measured after 24 h. Sucrose preference was defined as the ratio of sucrose solution consumed to total liquid intake during the test.

### Tail suspension test (TST)

This test was performed as described in the previous publication^74^ with minor modifications. Mice were suspended about 50 cm above the surface of an experimental table with adhesive tape, which was placed 1 cm from the tip of the tail to prevent the mouse from climbing upwards. Each mouse was tested for 6 min and the immobility time of the animal was measured in the last 5 min. The mouse was considered immobile when there were only small movements of the forefeet or no movement at all. Data acquisition and analysis were performed using a video tracking software (ANY-MAZE, Stoelting Co., Wood Dale, IL, USA).

### Sociability and social novelty test

This test was performed as described in the previous publications^75,76^ with minor modifications. The test apparatus consisted of a clear Plexiglas three-chamber box, with each chamber interconnected via sliding gates to allow free movement. The test consisted of three phases. Acclimation phase: prior to testing, each mouse was placed in the center chamber and allowed to explore all three chambers for 10 min to habituate to the apparatus; Sociability phase: following acclimation, the test mouse was returned to the center chamber, and the gates were closed. A novel stimulus mouse (sex- and age-matched C57BL/6J mouse with no prior contact) was enclosed in an inverted wire pencil cup in one of the side chambers, while an empty inverted wire pencil cup (serving as a novel object control) was placed in the opposite side chamber. The positions of the stimulus mouse and empty cup were counterbalanced across trials to avoid side bias. The gates were then reopened, and the test mouse was allowed to explore all three chambers for 10 min; Social novelty preference phase: Immediately after the sociability phase, the test mouse was again confined to the center chamber. The empty wire cup was replaced with a second novel stimulus mouse (sex- and age-matched, unfamiliar to the test mouse), while the original stimulus mouse remained in its chamber as a “familiar” social stimulus. The gates were reopened, and the test mouse was allowed to explore for an additional 10 min. All test sessions were video-recorded, and the duration of time the test mouse spent within 1 cm of each stimulus (novel object, familiar mouse, or novel mouse) was quantified using automated tracking software (ANY-maze 7.3). Reduced interaction time with novel conspecifics (versus objects or familiar mice) was interpreted as a deficit in social motivation, a behavioral correlation of depressive-like phenotypes.

### Western blot analysis

After *in vivo* or *in vitro* drug treatment, mouse hippocampi or hippocampal slices (prepared as described above) were dissected and pooled. Collected tissue was then homogenized in ice-cold RIPA buffer with 1× protease and phosphatase inhibitor cocktail. Homogenates (total fractions) were collected, and cell debris was removed by centrifugation at 13,000 rpm for 10 min at 4°C. Next, a bicinchoninic acid (BCA) protein assay kit (RG235625, Thermo) was used to determine the protein concentrations of the samples. Tissue samples were separated using 10% sodium dodecyl sulfate-polyacrylamide gel electrophoresis (SDS-PAGE), then transferred to polyvinylidene difluoride (PVDF) membranes and blocked with 5% non-fat milk in Tris-buffered saline (TBS) containing 0.05% Tween-20. PVDF membranes were incubated with antibodies as follows: mouse monoclonal anti-MOR-DOR heterodimer antibody (1:1000, Kerafast), or rabbit monoclonal anti-MOR antibody (1:1000, Proteintech), or rabbit monoclonal anti-DOR antibody (1:1000, Abcam), or rabbit monoclonal anti-GAPDH antibody (1:5000, Cell Signaling Technology, CST), or rabbit monoclonal anti-GluA1 antibody (1:8000, CST), or rabbit polyclonal anti-BDNF antibody (1:3000, CST), or rabbit monoclonal anti-β-actin antibody (1:20000, CST) at 4°C overnight following 1-h blocking. After washing with TBS-Tween, the membranes were incubated with horseradish peroxidase (HRP)-conjugated goat anti-mouse immunoglobulin G (IgG) (1:5000, Abcam) and HRP-conjugated goat anti-rabbit IgG (1:10000, Abcam). The immunoblot was developed using enhanced chemiluminescence (ECL) reagent (RJ240732, Thermo). Finally, blot quantification was performed using ImageQuant™ TL (IQTL) software (version 8), and densitometry values were normalized to the values obtained with the β-actin or GAPDH antibody. (For catalog numbers of antibodies, please see extended data table 3)

### Culture and co-culture of hippocampal neurons and astrocytes

*Culture of hippocampal neurons*: embryonic day 18 (E18) mouse embryos were collected from the uteri of euthanized pregnant mice. Following decapitation, embryonic brains were rapidly dissected and immediately immersed in ice-cold dissecting medium consisting of Dulbecco’s Modified Eagle Medium (DMEM, 12491023, Thermo) supplemented with 1% penicillin-streptomycin. For each independent experiment, bilateral hippocampal tissues were microdissected from 8–10 embryos on ice and pooled in the aforementioned dissecting medium. Hippocampal tissues were first digested with 0.25% trypsin-EDTA, followed by mechanical trituration using a fire-polished Pasteur pipette to achieve complete cellular dissociation.

The resulting cell suspension was filtered through a 70-μm cell strainer to remove undigested tissue clumps and debris. The filtered cell suspension was centrifuged at 130 × g for 10 min to pellet cells, which were then resuspended in complete DMEM containing 10% heat-inactivated fetal bovine serum (56°C for 30 min) and 1% penicillin-streptomycin. CultureOne™ supplement (#A3320201, Thermo) was added to the cell resuspension medium at this stage. Cells were seeded onto 13 mm round glass coverslips pre-coated with poly-D-lysine (PDL) in 24-well plates or directly onto the PDL-precoated bottom of NEST #723421 cell chambers. After an initial 5 h incubation at 37°C with 5% CO₂, the culture medium was fully replaced with neuronal-specific medium (21103049, Gibco) supplemented with 2% B27 supplement (A3582801, Gibco), 1% penicillin-streptomycin, and 1% GlutaMAX (#35050061, Gibco). To maintain primary neuronal cultures, half-volume medium changes were performed every 48 h, and neurons were cultured for 14 days to allow full maturation before subsequent experimental manipulations.

*Culture of astrocytes*: astrocytes were isolated from hippocampal tissues of E18 mouse embryos, using the same tissue source as the hippocampal neurons described above. Astrocytes were cultured in complete DMEM supplemented with 10% heat-inactivated fetal bovine serum and 1% penicillin-streptomycin, maintained at 37°C with 5% CO₂. Cultures were incubated for 9–14 days until astrocytes reached full confluence on the bottom of culture flasks. Astrocyte passaging was performed via enzymatic digestion: confluent astrocytes were rinsed three times with Hank’s Balanced Salt Solution (HBSS, 14175095, Gibco) to remove residual medium, then digested with 0.25% trypsin-EDTA. Digested astrocytes were gently triturated to form a single-cell suspension, which was then re-seeded onto the membrane surface of PDL-precoated NEST #723421 cell chambers for subsequent co-culture experiments.

*Co-culture of hippocampal neurons and astrocytes*: For the neuron-astrocyte indirect co-culture system, NEST #723421 cell chambers seeded with astrocytes were gently placed into wells of 6-well plates pre-seeded with mature hippocampal neurons. Extreme care was taken to ensure tight adherence of the cell chamber bottom to the 6-well plate well bottom, preventing chamber floating and air bubble formation between the chamber and plate—both of which would impair nutrient exchange across the chamber membrane. Neuronal-specific medium (supplemented with 2% B27, 1% penicillin-streptomycin, and 1% GlutaMAX) was used as the co-culture medium, added to both the lower well and upper cell chamber. The medium volume was adjusted to 2.5–3 mL in the lower 6-well plate well and 0.5–1 mL in the upper cell chamber, ensuring complete submersion of the chamber membrane to avoid membrane desiccation and subsequent cell death. Following establishment of the co-culture system, cells were treated with 10 μM HNK for 2 h, then processed for immunofluorescence staining and Western blot analysis as subsequent experimental assays.

### Immunofluorescence staining

On glass coverslips in 24-well plates, coverslips were fixed with 4% paraformaldehyde in 0.01 M PBS (pH 7.4) for 15 min at room temperature. Fixed cells were then permeabilized with 0.1% Triton X-100 in PBS for 30 min and blocked with 5% BSA in PBS for 1 h at room temperature to minimize nonspecific binding. Samples were then incubated with primary antibodies: rabbit anti-GluA1 (1:200, CST), mouse anti-Tuj1 (1:200, Biolegend), mouse anti-MOR-DOR heterodimer (1:200, Kerafast), rabbit anti-GFAP (1:500, Proteintech) overnight at 4°C in a humidified chamber. After three washes with 0.01 M PBS, samples were incubated with fluorophore-conjugated secondary antibodies: Alexa Fluor 488 goat anti mouse (1:1000, Invitrogen), Alexa Fluor 555 goat anti rabbit (1:1000, Invitrogen); Alexa Fluor 488 goat anti rabbit (1:1000, Invitrogen), Alexa Fluor 555 goat anti mouse (1:1000, Invitrogen) for 1 h at room temperature in the dark, followed by nuclear counterstaining with DAPI for 5 min. Coverslips were mounted with ProLong^TM^ Gold Antifade Mountant (Invitrogen) and imaged on a confocal microscope (Olympus FV3000) with 60× or 100× oil objectives. Fluorescence intensity and colocalization were quantified using NIH ImageJ software.

### *In vivo* two-photon calcium imaging

Mice were anesthetized with isoflurane (3% for induction and 1.5% for maintenance) and secured in a stereotaxic frame, with a thermostatic heating pad applied to maintain core body temperature at 37 °C. For dual-color cell-type-specific calcium labeling, two adeno-associated viral vectors were sequentially microinjected into the unilateral dorsal hippocampal CA1 region (AP: −2.0 mm, ML: −1.5 mm, DV: −1.4 mm, relative to Bregma). AAV2/5-GfaABC1D-GCaMP6f-WPRE-hGH poly A (300 nL, titer: 5.60 × 10^12^ vector genomes (v.g.)/mL) and AAV2/9-Syn-NES-SomaFRCaMPi (300 nL, titer: 5.45 × 10^12^ vector genomes (v.g.)/mL) were used to selectively label astrocytes (green fluorescence) and neurons (red fluorescence), respectively. Viral injection was performed at a constant rate of 30 nL/min, and the injection needle was retained in place for an additional 10 min following infusion completion to prevent viral backflow and ensure sufficient local diffusion.

After a 10-day postoperative adaptation recovery period, a 3-mm-diameter circular craniotomy was created over the right dorsal hippocampus under isoflurane anesthesia. The overlying cerebral cortex and corpus callosum were gently resected using a 26-gauge blunt needle connected to a vacuum pump to fully expose the intact dorsal hippocampal surface. Briefly, following craniotomy and dural removal, low-pressure suction (−50 to −60 kPa) was applied to gradually ablate superficial cortical and callosal tissues, with continuous artificial cerebrospinal fluid (ACSF) perfusion throughout the procedure to protect hippocampal tissue from thermal and mechanical damage. A custom stainless-steel cannula (3 mm diameter, 1.4 mm insertion depth) pre-assembled with a glass coverslip was implanted into the craniotomy using fine forceps. The cannula was firmly sealed to the skull with Vetbond™ tissue adhesive (3M), and a titanium head plate was anchored to the anterior cranial surface with dental cement for subsequent head fixation.

All mice were individually housed for at least two weeks post-surgery to allow full recovery and adequate viral expression before imaging acquisition. During formal imaging experiments, mice were anesthetized with low-dose isoflurane (3% induction, 1% maintenance) and head-fixed on a custom stage mounted on a vibration-isolation table to eliminate mechanical motion artifacts, with the body loosely restrained in a customized plastic holder. Calcium imaging was performed using an Olympus FVMPE-RS two-photon microscope equipped with a 25× water-immersion objective lens (NA = 1.05). Two distinct excitation wavelengths were applied for dual-channel imaging: 960 nm for GCaMP6f-labeled astrocytes and 1045 nm for SomaFRCaMPi-labeled neurons. Full-frame time-lapse images were acquired at 0.92 Hz for single-channel astrocytic recording and 0.31 Hz for simultaneous dual-color neuron–astrocyte imaging. For offline data analysis, raw calcium image sequences were first processed with motion correction in ImageJ to eliminate residual movement artifacts. Aligned image stacks were subsequently imported into the Suite2p pipeline for automated cell detection, region of interest (ROI) segmentation, and extraction of somatic fluorescence intensity traces (F). The baseline fluorescence (F0) of each individual cell was defined as the 20th percentile of its entire time-series fluorescence value. Relative calcium activity was calculated as the normalized fluorescence change: ΔF/F = (F − F0) / F0. To evaluate cumulative calcium activity intensity, the area under the curve (AUC) of ΔF/F traces was quantified via trapezoidal numerical integration across the entire recording duration.

All quantitative data are presented as mean ± standard error of the mean (SEM). Statistical analyses were conducted using GraphPad Prism software (version 10.1.2). Intergroup differences were assessed via independent samples Student’s t-test, and a p-value < 0.05 was defined as the threshold for statistical significance.

### Optogenetic manipulation

Mice were deeply anesthetized with 1.5% isoflurane. A viral vector was bilaterally injected into the CA1 region of the hippocampus in GFAP-Cre mice (coordinates relative to bregma: AP: −2.0 mm, ML: ±1.5 mm, DV: −1.5 mm; 300 nl of virus was delivered to the CA1 on each side). Subsequently, 2-mm optic fibers with a 1.25-mm ceramic ferrule (Inper Technology Co., Ltd., Hangzhou, China) were implanted targeting the same region (coordinates: AP: −2.0 mm, ML: ±1.5 mm, DV: −1.4 mm) and secured with dental cement. After surgery, animals were allowed to recover for 3 weeks. The following AAVs were used: AAV9-EF1a-DIO-EYFP (control, 2 × 10¹² vg·ml⁻¹), AAV9-EF1a-DIO-hChR2(H134R)-EYFP (9.12 × 10¹² vg·ml⁻¹), and AAV9-EF1a-DIO-eNpHR3.0-EYFP (2.07 × 10¹² vg·ml⁻¹).

For electrophysiological recordings, neurons were held at −70 mV. Only pyramidal neurons, identified by their morphological features and characteristic action potential patterns, were included in the statistical analysis. For patch-clamp experiments involving Channelrhodopsin-2 (ChR2), each hippocampal slice was used for only one optogenetic stimulation experiment, and only one neuron was patched per slice. If the recording became unstable after optogenetic stimulation, the slice was discarded. Spontaneous excitatory postsynaptic currents (sEPSCs) were recorded for an initial 5-min period without light stimulation. This was followed by delivery of 460-nm light (20 Hz, 5-ms pulse width) to the slice for 5 min. Next, 10 μM HNK was added to the bath solution, and the light was turned off for 5 min. Finally, light stimulation resumed, and sEPSCs were recorded for an additional 5 min. For experiments involving Halorhodopsin (NpHR), 560-nm continuous light was delivered to the hippocampal slice; all other procedures remained identical. Light stimulation was delivered using a light-stimulation system (#MG-120, Mshot Optical Instrument Co., Ltd., Guangzhou, China).

### Bimolecular fluorescence complementarity (BiFC) analysis

The coding sequences of *Oprm1* (NM_000914.5) *and Oprd1* (NM_000911.4) were cloned into the N-terminal fragment (VN, 1-172 aa) and C-terminal fragment (VC, 155-239 aa) expression vectors (pcDNA3.1+) of mVenus, respectively, to construct the fusion plasmids *Oprm1*-VN172 and *Oprd1*-VC155. Negative controls included non-interacting protein pairs (VN-C+VC-D), single vector transfection (VN-A+empty VC), and key site mutants (VN-A-mut+VC-B). All plasmids were validated by sequencing and extracted using the EndoFree Plasmid Maxi Kit (DC202, Vazyme). After inoculating CHO cells into a 6-well plate (density of 60-70%), Lipo8000™ transfection reagent was used to co-transfect *Oprm1*-VN172 and *Oprd1*-VC155 (molar ratio 1:1, total plasmid 2 μg/well). After transfection for 6 h, the culture medium was replaced, and the culture was continued for 24-48 h. Transfected cells were digested with trypsin and seeded into confocal dishes. Time-series fluorescence images of live cells were captured using a live-cell ultra-high-resolution confocal imaging system (SpinSR disk confocal, 60× oil objective) with a 514 nm argon laser. Imaging parameters were fixed to avoid photobleaching. The real-time fluorescence intensity at the cytoplasmic membrane surface was quantified from time-series images using ImageJ.

### Dual-target PAINT-MINFLUX imaging

#### Sample preparation

CHO cells were co-transfected with MOR-EGFP and DOR-ALFA plasmids using Lipofectamine 3000 (#L3000015, Thermo). At 24 h post-transfection, cells were detached, resuspended, and seeded onto 18 mm circular glass coverslips. They were then divided into two groups: an experimental group and a peptide-treated group. The peptide-treated group was incubated with PEP (50 nM). After an additional 24 h, both groups were subjected to the following treatments for 5 minutes at 37°C: Control (PBS), HNK (agonist, 10 μM), or CYM (agonist, 1 μM). Following treatment, cells were immediately fixed with 4% paraformaldehyde (PFA) for 10 minutes. Subsequently, cells were washed three times with PBS containing 100 mM glycine. Cells were then permeabilized and blocked for ≥30 min at room temperature in a blocking buffer containing 0.3% Triton X-100, 3% BSA and 10% donkey serum. For dual-color labeling, Anti-GFP and anti-ALFA nanobodies conjugated to distinct docking strands (Massive Photonics) were diluted 1:100 in antibody incubation buffer and applied for 1 h at room temperature. After three washes with PBS, cells were post-fixed with 4% PFA for 5 min. To serve as fiducial markers for drift correction, samples were incubated with 200 μL gold nanoparticles (250 nm colloidal gold; BBI Solutions, Cardiff, Wales, UK) for 10 min. This was followed by multiple PBS washes to remove unbound particles. Cy3b-conjugated DNA-PAINT imager strands (Massive Photonics) were diluted in imaging buffer to 200-500 pM. Prepared samples were mounted in a custom chamber containing the respective imager solution. Data acquisition was completed within 2 days of labeling.

#### Imaging

Imaging was performed on a commercial Abberior MINFLUX system built on an inverted IX83 frame (Olympus) with a 100× UPlanXApo objective (Olympus) and controlled by Imspector (Abberior Instruments, Göttingen, Lower Saxony, Germany). The excitation beam pattern and pinhole position were calibrated daily using fluorescent nanoparticles (Abberior Nanoparticles; Gold 150 nm, 2C Fluor 120 nm). Cells co-expressing GFP-MOR and ALFA-DOR were identified via488 nm confocal scanning, and the focal plane was set at the coverslip-cell interface. Transient binding of Cy3b imagers was verified with 560 nm confocal scanning. Each field contained at least three gold fiducials and was stabilized in real time by the microscope’s active feedback system, achieving < 1 nm precision in x/y. Regions of interest (ROIs) were recorded using a standard 2D MINFLUX sequence with the pinhole set to 0.83 arbitrary units (A.U.). Following the first imaging round (e.g., for MOR), the chamber solution was exchanged via a custom perfusion system until the capture rate decayed to zero, indicating complete removal of the first imager. The second, orthogonal imager (for DOR) was then introduced by perfusion, and the second PAINT-MINFLUX imaging round commenced. MINFLUX acquisition continued uninterrupted throughout, with continuous active-feedback stabilization, obviating post hoc drift correction. Both imaging rounds plus solution exchange were typically completed within about 3 h.

#### Data analysis

Localization coordinates from converged MINFLUX events were exported from Imspector as .mat files. All subsequent post-processing, filtering, and visualization were performed in MATLAB using custom scripts. Applied filters included: (1) a center frequency ratio (CFR) threshold of 0.8 (applied during acquisition); (2) an effective frequency at offset (EFO, kHz) to exclude potential multiple, closely spaced binding events; and (3) an aggregation step where localizations originating from the same binding event (coordinates sharing the same TID) were grouped, retaining only events with > 5 repeats and replacing them by their mean coordinate. To assign localizations to their respective targets, the manually recorded time of second-imager introduction served as the segmentation boundary: all localizations before this time were assigned to molecule A (e.g., MOR), and those after to molecule B (e.g., DOR). Considering the localization precision (∼3 nm), the size of the nanobody (∼5 nm), and the size and flexibility of the receptor (∼10 nm), a 20 nm center-to-center distance threshold was used to define potential dimerized pairs. The probability of random colocalization events being misidentified as dimers was corrected using a coupled-probability algorithm^61^. Briefly, Monte Carlo simulations generated random points within ranges defined by inter-molecular distances and localization precision; the proximity probability was computed as the fraction of simulated points falling within the 20 nm threshold. Using a bipartite-graph formulation with a non-forced greedy matching, the optimal number of pairings was identified. To estimate the background level of random pairings due to molecular crowding, an iterative approach based on iMEC was employed. In this process, the coordinates of molecule B were randomly reassigned and re-matched to molecule A coordinates over multiple iterations until convergence. The final set of bona fide dimers was obtained by subtracting these estimated background matches from the initial optimal pairings and then re-optimizing the pairings.

### Atomistic classical molecular dynamics simulation

Given that the heterodimeric structure of the MOR and DOR has not been experimentally established, the MOR-DOR heterodimer complex was constructed using AlphaFold3 (**Extended Data Fig. 7**)^77^. The binding site of HNK to the heterodimer was also predicted using AlphaFold3. This prediction was further validated through atomistic classical molecular dynamics (MD) simulations of the heterodimer in the presence of HNK (**Extended Data Fig. 8**). The transmembrane structure of the MOR-DOR heterodimers was prepared using CHARMM-GUI (https://www.charmm-gui.org/)^78^. Details of the molecular system are summarized in Table 2.

The MD simulation was executed using the GROMACS 2020.3 software package^79^, coupled with the Amber19SB force field^80^. HNK topology was prepared with the AmberTools package, and its bonded and nonbonded interaction parameters were derived from the General Amber Force Field (GAFF)^81^. Partial charges for the molecule were calculated using restrained electrostatic potential (RESP) fitting, based on Hartree–Fock calculations performed with the 6-31G* basis set^82^. The simulation system was solvated with TIP3P water molecules, and sodium (Na⁺) and chloride (Cl⁻) ions were introduced to neutralize the system charge and emulate a physiological ionic strength of 0.15 M NaCl.

Simulations were performed under conditions of 310 K temperature and 1 bar pressure. Temperature coupling was achieved using the velocity-rescaling algorithm^83^, while pressure regulation employed the Parrinello–Rahman barostat^84^. Long-range electrostatics were handled via the particle mesh Ewald (PME) method, applying a real-space cutoff of 1.4 nm. The same cut-off distance was used for van der Waals interactions. Periodic boundary conditions were applied along all axes to maintain consistency across the simulation box. The interaction energies between specific regions of the MOR-DOR heterodimers and ketamine were calculated using the MM/PBSA (Molecular Mechanics/Poisson–Boltzmann Surface Area) method^85^.

### Site-directed mutagenesis and plasmid construction

The specific primers for point mutation were designed according to the mutation sites. The melting temperature was optimized to 60-65°C, and the homologous region at the flanks was 20-25 base pairs (bp). PCR amplification was performed using Phanta Max Master Mix (#P515, Vazyme Biotech Co., Ltd., Nanjing, Jiangsu, China) high-fidelity DNA polymerase in a 50 μL reaction containing 10 ng of wild-type plasmid template. The PCR product was digested with 20 U DMT Enzyme (#GD111-01, TransGen Biotech, Inc., Beijing, China) at 37°C for 1 h to eliminate methylated parental DNA. The product was then purified using a DNA cleanup and concentration kit (#DC301, Vazyme) and was transformed into *E. coli* DH5α competent cells (TransGen Biotech) by heat shock. Transformed colonies were screened on LB agar plates containing 100 μg·mL^-1^ ampicillin, and colony PCR was performed on individual colonies for preliminary validation. Bidirectional sequencing (conducted by GENEWIZ) was performed on plasmids exhibiting the correct restriction pattern using the CMV forward primers. The resulting sequences were aligned with the reference sequence using SnapGene software to verify the accuracy of the mutation. Validated plasmids were amplified in LB medium and purified using the EndoFree Plasmid Maxi Kit (DC202, Vazyme), with their concentrations confirmed by NanoDrop (A260/A280 ≥ 1.8). Wild-type plasmids and template-free PCR reactions served as negative controls.

### Co-immunoprecipitation

Upon reaching 70-80% confluence, cultured HEK293T cells (obtained from the American Type Culture Collection, ATCC) were co-transfected with pcDNA3.1-MOR-3×Flag and pcDNA3.1-DOR-3×HA plasmids. Forty-eight hours post-transfection, cells were washed twice with ice-cold PBS and lysed in ice-cold cell lysis buffer (#P0013J, Beyotime Biotechnology Co., Ltd., Shanghai, China) supplemented with protease and phosphatase inhibitor cocktails (Roche cOmplete™) for 30 min on ice. Lysates were centrifuged at 12,000× g for 15 min at 4°C, and the supernatants were collected. Protein concentration was determined using the BCA Protein Assay Kit (RG235625, Thermo) with BSA as the standard. 1-2 μg of anti-DYKDDDDK tag recombinant antibody (#80801-2-RR, Proteintech Group, Inc., Wuhan, China) was incubated with 20 μL of Protein A/G magnetic beads (#HY-K0202, MCE) in IP buffer for 2 h at 4°C to form antibody-bead complexes. A negative control group was set up using isotype-matched IgG (A7028, Beyotime) instead of the primary antibody. Equal amounts (500-1000 μg) of protein lysates were incubated with antibody-bead complexes overnight at 4°C with gentle rotation. Following incubation, beads were washed four times with ice-cold lysis buffer, and centrifuged at 1,000× g for 5 min each time at 4°C. Bound proteins were eluted by adding 2× SDS-PAGE loading buffer containing DTT and heating at 95°C for 10 min. Eluted samples were separated by SDS-PAGE and transferred to a PVDF membrane (#ISEQ00010, MilliporeSigma). Membranes were blocked with 5% non-fat milk, probed with primary antibodies as follow: anti-HA tag (#M20003S, Abmart Co., Ltd., Shanghai, China); anti-Flag tag (#M20008S, Abmart), followed by HRP-conjugated secondary antibodies, and visualized using ECL substrate (#P10300, NCM Biotech Co., Ltd., Suzhou, China).

### Statistical information

All statistical analyses were conducted using GraphPad Prism version 10. Prior to the statistical analysis, the normality and homogeneity of variances of the data were assessed. Normality was evaluated using the Shapiro-Wilk test, and homogeneity of variances was confirmed using the Brown-Forsythe test. Both normality and homogeneity of variances were confirmed in this study. For the analysis of Western blot, electrophysiology, and behavior tests, data were presented as the mean ± standard error of the mean (SEM). Test mice that failed to complete the current behavioral assessment were excluded from subsequent testing based on predefined exclusion criteria; these data were analyzed by two-tailed Student’s t-test or one-way analysis of variance (ANOVA) with Fisher’s least significant difference (LSD) post hoc test, based on the data types. In electrophysiological experiments, fEPSP or EPSC slope values were averaged and quantified over a 10-min period preceding application of the substance (control) and a 10-min period at the end of the substance application. A P value of less than 0.05 was considered statistically significant.

## Data availability

All data are available in the manuscript or the supplementary materials.

## Acknowledgements

We thank Dr. Yan Liu and Dr. Lei Guo for advice on immunohistochemical staining; Dr. Liu Zheng, Qing Zhang, and Qi Wang for technical support; Xiao-Qin Huang for assistant on optogenetic experiments; We acknowledge the use of imaging facilities at the Multimodality Imaging Center of the Institute of Artificial Intelligence of Hefei Comprehensive National Science Center. This work was supported by grants from the National Natural Science Foundation of China (82471548 to X. C.; 82471538 to C. W.; 3241007 and T2596040 to A.-H. T.), Natural Science Funds for Distinguished Young Scholar of Zhejiang (LR20H090001 to C. W.), Brain Science and Brain-like Intelligence Technology-National Science and Technology Major Project (2021ZD0202500 to A.-H. T.), the Open Fund of Anhui Province Key Laboratory of Biomedical Imaging and Intelligent Processing (24YGXT002 to A.-H. T.), Public Welfare Scientific Research Foundation of Wenzhou (Y2023064 to X. C.), Ningbo Youth Leading Talent Project (2025QL074 to Y.-X. S.), the Yunnan Revitalization Talent Support Program Yunling Scholar Project (M. L.).

## Author contributions

Y.-X. Liang, X.-X. Li, M.-L. Yang, C. Xu, Y. Chen, H.-Y. Bai and S. Yang performed behavioural experiments, biochemistry, and immunohistochemistry experiments; Y.-J. Li, X.-X. Li, and Y.-X. Liang conducted viral injection, cell cultures, plasmid construction, BiFC, and co-immunoprecipitation; L.-J. Wang, C. Xu, R. Thapa, S. Yan, H.-Q. Zhang, Z.-C. He, Z.-X. Fang performed electrophysiological experiments; L.-J. Wang performed optogenetic experiments; A.-H. Tang, X.-Z. Guo, and J.-X. Mu designed and performed PAINT-MINFLUX experiments. C. Wang and Y.-X. Sun designed and performed atomistic classical molecular dynamics simulation; J.-Z. Su, Q. Zhang, J. Xu, Z. Liu conducted data analysis; X. Cai designed the study, and X. Cai wrote the manuscript with assistance of Q. Zhou, C. Wang, A.-H. Tang, Y.-X. Sun, M. Li, and W.-J. Geng, and D.-W. Xu.

## Correspondence and requests for materials

Xiang Cai

## Reference

1. Berman RM, Cappiello A, Anand A, et al. Antidepressant effects of ketamine in depressed patients. Biol Psychiatry. Feb 15 2000;47(4):351–4. doi:10.1016/s0006-3223(99)00230-9

2. Li N, Lee B, Liu RJ, et al. mTOR-dependent synapse formation underlies the rapid antidepressant effects of NMDA antagonists. Science. Aug 20 2010;329(5994):959–64. doi:10.1126/science.1190287

3. McIntyre RS, Alsuwaidan M, Baune BT, et al. Treatment-resistant depression: definition, prevalence, detection, management, and investigational interventions. World Psychiatry. Oct 2023;22(3):394–412. doi:10.1002/wps.21120

4. Autry AE, Adachi M, Nosyreva E, et al. NMDA receptor blockade at rest triggers rapid behavioural antidepressant responses. Nature. Jun 15 2011;475(7354):91–5. doi:10.1038/nature10130

5. Zanos P, Moaddel R, Morris PJ, et al. Ketamine and Ketamine Metabolite Pharmacology: Insights into Therapeutic Mechanisms. Pharmacol Rev. Jul 2018;70(3):621–660. doi:10.1124/pr.117.015198

6. Zhang K, Yamaki VN, Wei Z, Zheng Y, Cai X. Differential regulation of GluA1 expression by ketamine and memantine. Behav Brain Res. Jan 1 2017;316:152–159. doi:10.1016/j.bbr.2016.09.002

7. Zanos P, Moaddel R, Morris PJ, et al. NMDAR inhibition-independent antidepressant actions of ketamine metabolites. Nature. May 26 2016;533(7604):481–6. doi:10.1038/nature17998

8. Raja SM, Guptill JT, Mack M, et al. A Phase 1 Assessment of the Safety, Tolerability, Pharmacokinetics and Pharmacodynamics of (2R,6R)-Hydroxynorketamine in Healthy Volunteers. Clin Pharmacol Ther. Nov 2024;116(5):1314–1324. doi:10.1002/cpt.3391

9. Lumsden EW, Troppoli TA, Myers SJ, et al. Antidepressant-relevant concentrations of the ketamine metabolite (2R,6R)-hydroxynorketamine do not block NMDA receptor function. Proc Natl Acad Sci U S A. Mar 12 2019;116(11):5160–5169. doi:10.1073/pnas.1816071116

10. Fukumoto K, Fogaça MV, Liu RJ, et al. Activity-dependent brain-derived neurotrophic factor signaling is required for the antidepressant actions of (2R,6R)-hydroxynorketamine. Proc Natl Acad Sci U S A. Jan 2 2019;116(1):297–302. doi:10.1073/pnas.1814709116

11. Wang H, He Y, Tang J, et al. (2R, 6R)-hydroxynorketamine ameliorates PTSD-like behaviors during the reconsolidation phase of fear memory in rats by modulating the VGF/BDNF/GluA1 signaling pathway in the hippocampus. Behav Brain Res. Jan 5 2025;476:115273. doi:10.1016/j.bbr.2024.115273

12. Thompson SM, Kallarackal AJ, Kvarta MD, Van Dyke AM, LeGates TA, Cai X. An excitatory synapse hypothesis of depression. Trends Neurosci. May 2015;38(5):279–94. doi:10.1016/j.tins.2015.03.003

13. Munguba H, Arefin A, Hasegawa R, et al. Mechanism-guided identification of antidepressant G protein-coupled receptor drug targets. Cell. Apr 30 2026;189(9):2612–2632 e24. doi:10.1016/j.cell.2026.04.006

14. Aguilar-Valles A, De Gregorio D, Matta-Camacho E, et al. Antidepressant actions of ketamine engage cell-specific translation via eIF4E. Nature. Feb 2021;590(7845):315–319. doi:10.1038/s41586-020-03047-0

15. Zhao YF, Verkhratsky A, Tang Y, Illes P. Astrocytes and major depression: The purinergic avenue. Neuropharmacology. Dec 1 2022;220:109252. doi:10.1016/j.neuropharm.2022.109252

16. Peng L, Verkhratsky A, Gu L, Li B. Targeting astrocytes in major depression. Expert Rev Neurother. 2015;15(11):1299–306. doi:10.1586/14737175.2015.1095094

17. Wang Q, Jie W, Liu JH, Yang JM, Gao TM. An astroglial basis of major depressive disorder? An overview. Glia. Aug 2017;65(8):1227–1250. doi:10.1002/glia.23143

18. Xin Q, Wang J, Zheng J, et al. Neuron-astrocyte coupling in lateral habenula mediates depressive-like behaviors. Cell. Jun 12 2025;188(12):3291–3309 e24. doi:10.1016/j.cell.2025.04.010

19. Cornell-Bell AH, Finkbeiner SM, Cooper MS, Smith SJ. Glutamate induces calcium waves in cultured astrocytes: long-range glial signaling. Science. Jan 26 1990;247(4941):470–3. doi:10.1126/science.1967852

20. Cahill MK, Collard M, Tse V, et al. Network-level encoding of local neurotransmitters in cortical astrocytes. Nature. May 2024;629(8010):146–153. doi:10.1038/s41586-024-07311-5

21. Volterra A, Liaudet N, Savtchouk I. Astrocyte Ca(2)(+) signalling: an unexpected complexity. Nat Rev Neurosci. May 2014;15(5):327–35. doi:10.1038/nrn3725

22. Zhang JM, Wang HK, Ye CQ, et al. ATP released by astrocytes mediates glutamatergic activity-dependent heterosynaptic suppression. Neuron. Dec 4 2003;40(5):971–82. doi:10.1016/s0896-6273(03)00717-7

23. Nedergaard M. Direct signaling from astrocytes to neurons in cultures of mammalian brain cells. Science. Mar 25 1994;263(5154):1768–71. doi:10.1126/science.8134839

24. Sun W, Liu Z, Jiang X, et al. Spatial transcriptomics reveal neuron-astrocyte synergy in long-term memory. Nature. Mar 2024;627(8003):374–381. doi:10.1038/s41586-023-07011-6

25. Albini M, Krawczun-Rygmaczewska A, Cesca F. Astrocytes and brain-derived neurotrophic factor (BDNF). Neurosci Res. Dec 2023;197:42–51. doi:10.1016/j.neures.2023.02.001

26. Tewari BP, Woo AM, Prim CE, et al. Astrocytes require perineuronal nets to maintain synaptic homeostasis in mice. Nat Neurosci. Aug 2024;27(8):1475–1488. doi:10.1038/s41593-024-01714-3

27. Subba R, Ellappan S, Banerjee S, Mondal AC. Astroglia and depression: A Gliocentric perspective from rodent models to therapeutic insights. Prog Neuropsychopharmacol Biol Psychiatry. Mar 20 2026;145:111627. doi:10.1016/j.pnpbp.2026.111627

28. Chi D, Zhang K, Zhang J, et al. Astrocytic pleiotrophin deficiency in the prefrontal cortex contributes to stress-induced depressive-like responses in male mice. Nat Commun. Mar 14 2025;16(1):2528. doi:10.1038/s41467-025-57924-1

29. Cui Y, Yang Y, Ni Z, et al. Astroglial Kir4.1 in the lateral habenula drives neuronal bursts in depression. Nature. Feb 14 2018;554(7692):323–327. doi:10.1038/nature25752

30. Berrocoso E, Sanchez-Blazquez P, Garzon J, Mico JA. Opiates as antidepressants. Curr Pharm Des. 2009;15(14):1612-22. doi:10.2174/138161209788168100

31. Peciña M, Karp JF, Mathew S, Todtenkopf MS, Ehrich EW, Zubieta JK. Endogenous opioid system dysregulation in depression: implications for new therapeutic approaches. Mol Psychiatry. Apr 2019;24(4):576–587. doi:10.1038/s41380-018-0117-2

32. Browne CA, Jacobson ML, Lucki I. Novel Targets to Treat Depression: Opioid-Based Therapeutics. Harv Rev Psychiatry. Jan/Feb 2020;28(1):40–59. doi:10.1097/HRP.0000000000000242

33. Mathis VP, Ehrlich AT, Darcq E. The neural circuits and signalling pathways of opioid use disorder. Nat Rev Neurosci. Dec 2025;26(12):778–797. doi:10.1038/s41583-025-00982-7

34. Spodnick MB, McElderry SC, Diaz MR. Opioid receptor signaling throughout ontogeny: Shaping neural and behavioral trajectories. Neurosci Biobehav Rev. Mar 2025;170:106033. doi:10.1016/j.neubiorev.2025.106033

35. Zhang L, Zhang JT, Hang L, Liu T. Mu Opioid Receptor Heterodimers Emerge as Novel Therapeutic Targets: Recent Progress and Future Perspective. Front Pharmacol. 2020;11:1078. doi:10.3389/fphar.2020.01078

36. Jordan BA, Devi LA. G-protein-coupled receptor heterodimerization modulates receptor function. Nature. Jun 17 1999;399(6737):697–700. doi:10.1038/21441

37. Costantino CM, Gomes I, Stockton SD, Lim MP, Devi LA. Opioid receptor heteromers in analgesia. Expert Rev Mol Med. Apr 10 2012;14:e9. doi:10.1017/erm.2012.5

38. Zhang K, Xu T, Yuan Z, et al. Essential roles of AMPA receptor GluA1 phosphorylation and presynaptic HCN channels in fast-acting antidepressant responses of ketamine. Sci Signal. Dec 13 2016;9(458):ra123. doi:10.1126/scisignal.aai7884

39. Riggs LM, Aracava Y, Zanos P, et al. (2R,6R)-hydroxynorketamine rapidly potentiates hippocampal glutamatergic transmission through a synapse-specific presynaptic mechanism. Neuropsychopharmacology. Jan 2020;45(2):426–436. doi:10.1038/s41386-019-0443-3

40. Riggs LM, Thompson SM, Gould TD. (2R,6R)-hydroxynorketamine rapidly potentiates optically-evoked Schaffer collateral synaptic activity. Neuropharmacology. Aug 15 2022;214:109153. doi:10.1016/j.neuropharm.2022.109153

41. Williams NR, Heifets BD, Blasey C, et al. Attenuation of Antidepressant Effects of Ketamine by Opioid Receptor Antagonism. Am J Psychiatry. Dec 1 2018;175(12):1205–1215. doi:10.1176/appi.ajp.2018.18020138

42. Williams NR, Heifets BD, Bentzley BS, et al. Attenuation of antidepressant and antisuicidal effects of ketamine by opioid receptor antagonism. Mol Psychiatry. Dec 2019;24(12):1779–1786. doi:10.1038/s41380-019-0503-4

43. Zhang F, Hillhouse TM, Anderson PM, et al. Opioid receptor system contributes to the acute and sustained antidepressant-like effects, but not the hyperactivity motor effects of ketamine in mice. Pharmacol Biochem Behav. Sep 2021;208:173228. doi:10.1016/j.pbb.2021.173228

44. Klein ME, Chandra J, Sheriff S, Malinow R. Opioid system is necessary but not sufficient for antidepressive actions of ketamine in rodents. Proc Natl Acad Sci U S A. Feb 4 2020;117(5):2656–2662. doi:10.1073/pnas.1916570117

45. McQuiston AR, Saggau P. Mu-opioid receptors facilitate the propagation of excitatory activity in rat hippocampal area CA1 by disinhibition of all anatomical layers. J Neurophysiol. Sep 2003;90(3):1936–48. doi:10.1152/jn.01150.2002

46. Nam MH, Won W, Han KS, Lee CJ. Signaling mechanisms of mu-opioid receptor (MOR) in the hippocampus: disinhibition versus astrocytic glutamate regulation. Cell Mol Life Sci. Jan 2021;78(2):415–426. doi:10.1007/s00018-020-03595-8

47. Saitoh A, Yamada M. Antidepressant-like Effects of delta Opioid Receptor Agonists in Animal Models. Curr Neuropharmacol. Sep 2012;10(3):231–8. doi:10.2174/157015912803217314

48. Jutkiewicz EM. The antidepressant-like effects of delta-opioid receptor agonists. Mol Interv. Jun 2006;6(3):162–9. doi:10.1124/mi.6.3.7

49. Yekkirala AS, Kalyuzhny AE, Portoghese PS. Standard opioid agonists activate heteromeric opioid receptors: evidence for morphine and [d-Ala(2)-MePhe(4)-Glyol(5)]enkephalin as selective mu-delta agonists. ACS Chem Neurosci. Feb 17 2010;1(2):146–54. doi:10.1021/cn9000236

50. Gomes I, Jordan BA, Gupta A, Trapaidze N, Nagy V, Devi LA. Heterodimerization of mu and delta opioid receptors: A role in opiate synergy. J Neurosci. Nov 15 2000;20(22):RC110. doi:10.1523/JNEUROSCI.20-22-j0007.2000

51. Wu B, Hand W, Alexov E. Opioid Addiction and Opioid Receptor Dimerization: Structural Modeling of the OPRD1 and OPRM1 Heterodimer and Its Signaling Pathways. Int J Mol Sci. Sep 24 2021;22(19)doi:10.3390/ijms221910290

52. Nam MH, Han KS, Lee J, et al. Expression of micro-Opioid Receptor in CA1 Hippocampal Astrocytes. Exp Neurobiol. Apr 2018;27(2):120–128. doi:10.5607/en.2018.27.2.120

53. Svoboda KR, Adams CE, Lupica CR. Opioid receptor subtype expression defines morphologically distinct classes of hippocampal interneurons. J Neurosci. Jan 1 1999;19(1):85–95. doi:10.1523/jneurosci.19-01-00085.1999

54. Drake CT, Milner TA. Mu opioid receptors are in somatodendritic and axonal compartments of GABAergic neurons in rat hippocampal formation. Brain Res. Dec 4 1999;849(1-2):203–15. doi:10.1016/s0006-8993(99)01910-1

55. Drake CT, Milner TA. Mu opioid receptors are in discrete hippocampal interneuron subpopulations. Hippocampus. 2002;12(2):119–36. doi:10.1002/hipo.1107

56. Jiang C, Wang X, Le Q, et al. Morphine coordinates SST and PV interneurons in the prelimbic cortex to disinhibit pyramidal neurons and enhance reward. Mol Psychiatry. Apr 2021;26(4):1178–1193. doi:10.1038/s41380-019-0480-7

57. Nam MH, Han KS, Lee J, et al. Activation of Astrocytic mu-Opioid Receptor Causes Conditioned Place Preference. Cell Rep. Jul 30 2019;28(5):1154–1166 e5. doi:10.1016/j.celrep.2019.06.071

58. Rezai X, Faget L, Bednarek E, Schwab Y, Kieffer BL, Massotte D. Mouse delta opioid receptors are located on presynaptic afferents to hippocampal pyramidal cells. Cell Mol Neurobiol. May 2012;32(4):509–16. doi:10.1007/s10571-011-9791-1

59. Erbs E, Faget L, Scherrer G, et al. Distribution of delta opioid receptor-expressing neurons in the mouse hippocampus. Neuroscience. Sep 27 2012;221:203–13. doi:10.1016/j.neuroscience.2012.06.023

60. Commons KG, Milner TA. Localization of delta opioid receptor immunoreactivity in interneurons and pyramidal cells in the rat hippocampus. J Comp Neurol. May 12 1997;381(3):373–87.

61. Deguchi T, Iwanski MK, Schentarra EM, et al. Direct observation of motor protein stepping in living cells using MINFLUX. Science. Mar 10 2023;379(6636):1010–1015. doi:10.1126/science.ade2676

62. Wirth JO, Scheiderer L, Engelhardt T, Engelhardt J, Matthias J, Hell SW. MINFLUX dissects the unimpeded walking of kinesin-1. Science. Mar 10 2023;379(6636):1004–1010. doi:10.1126/science.ade2650

63. Balzarotti F, Eilers Y, Gwosch KC, et al. Nanometer resolution imaging and tracking of fluorescent molecules with minimal photon fluxes. Science. Feb 10 2017;355(6325):606–612. doi:10.1126/science.aak9913

64. Steen PR, Unterauer EM, Masullo LA, et al. The DNA-PAINT palette: a comprehensive performance analysis of fluorescent dyes. Nat Methods. Sep 2024;21(9):1755–1762. doi:10.1038/s41592-024-02374-8

65. Liu RJ, Lee FS, Li XY, Bambico F, Duman RS, Aghajanian GK. Brain-derived neurotrophic factor Val66Met allele impairs basal and ketamine-stimulated synaptogenesis in prefrontal cortex. Biol Psychiatry. Jun 1 2012;71(11):996–1005. doi:10.1016/j.biopsych.2011.09.030

66. Hashimoto K. Molecular mechanisms of the rapid-acting and long-lasting antidepressant actions of (R)-ketamine. Biochem Pharmacol. Jul 2020;177:113935. doi:10.1016/j.bcp.2020.113935

67. Zanos P, Gould TD. Mechanisms of ketamine action as an antidepressant. Mol Psychiatry. Apr 2018;23(4):801–811. doi:10.1038/mp.2017.255

68. Su T, Lu Y, Fu C, Geng Y, Chen Y. GluN2A mediates ketamine-induced rapid antidepressant-like responses. Nat Neurosci. Oct 2023;26(10):1751–1761. doi:10.1038/s41593-023-01436-y

69. Yue C, Wang N, Zhai H, et al. Adenosine signalling drives antidepressant actions of ketamine and ECT. Nature. Jan 2026;649(8096):423–431. doi:10.1038/s41586-025-09755-9

70. Yang Y, Cui Y, Sang K, et al. Ketamine blocks bursting in the lateral habenula to rapidly relieve depression. Nature. Feb 14 2018;554(7692):317–322. doi:10.1038/nature25509

71. Cao X, Li LP, Wang Q, et al. Astrocyte-derived ATP modulates depressive-like behaviors. Nat Med. Jun 2013;19(6):773–7. doi:10.1038/nm.3162

72. Cai X, Liang CW, Muralidharan S, Kao JP, Tang CM, Thompson SM. Unique roles of SK and Kv4.2 potassium channels in dendritic integration. Neuron. Oct 14 2004;44(2):351–64. doi:10.1016/j.neuron.2004.09.026

73. Cai X, Kallarackal AJ, Kvarta MD, et al. Local potentiation of excitatory synapses by serotonin and its alteration in rodent models of depression. Nat Neurosci. Apr 2013;16(4):464–72. doi:10.1038/nn.3355

74. Cui M, Ji R, Song L, et al. Neuronal and Molecular Mechanisms Underlying Chronic Pain and Depression Comorbidity in the Paraventricular Thalamus. J Neurosci. Mar 27 2024;44(13)doi:10.1523/JNEUROSCI.1752-23.2024

75. Zain MA, Pandy V, Majeed ABA, Wong WF, Mohamed Z. Chronic restraint stress impairs sociability but not social recognition and spatial memoryin C57BL/6J mice. Exp Anim. Feb 26 2019;68(1):113–124. doi:10.1538/expanim.18-0078

76. Carter MD, Shah CR, Muller CL, Crawley JN, Carneiro AM, Veenstra-VanderWeele J. Absence of preference for social novelty and increased grooming in integrin beta3 knockout mice: initial studies and future directions. Autism Res. Feb 2011;4(1):57–67. doi:10.1002/aur.180

77. Abramson J, Adler J, Dunger J, et al. Accurate structure prediction of biomolecular interactions with AlphaFold 3. Nature. Jun 2024;630(8016):493–500. doi:10.1038/s41586-024-07487-w

78. Jo S, Kim T, Iyer VG, Im W. CHARMM-GUI: a web-based graphical user interface for CHARMM. J Comput Chem. Aug 2008;29(11):1859–65. doi:10.1002/jcc.20945

79. Mark James Abrahama, Teemu Murtolad, Roland Schulzbc, Szilard Palla, Jeremy C. Smithb, Berk Hessa, Erik Lindahla,. GROMACS: High performance molecular simulations through multi-level parallelism from laptops to supercomputers. Software X. 2015;1-2(2015):19–25. doi:10.1016/j.softx.2015.06.001

80. Tian C, Kasavajhala K, Belfon KAA, et al. ff19SB: Amino-Acid-Specific Protein Backbone Parameters Trained against Quantum Mechanics Energy Surfaces in Solution. J Chem Theory Comput. Jan 14 2020;16(1):528–552. doi:10.1021/acs.jctc.9b00591

81. Case DA, Cheatham TE, 3rd, Darden T, et al. The Amber biomolecular simulation programs. J Comput Chem. Dec 2005;26(16):1668–88. doi:10.1002/jcc.20290

82. Barca GMJ, Bertoni C, Carrington L, et al. Recent developments in the general atomic and molecular electronic structure system. J Chem Phys. Apr 21 2020;152(15):154102. doi:10.1063/5.0005188

83. Bussi G, Donadio D, Parrinello M. Canonical sampling through velocity rescaling. J Chem Phys. Jan 7 2007;126(1):014101. doi:10.1063/1.2408420

84. Aoki KM, Yonezawa F. Constant-pressure molecular-dynamics simulations of the crystal-smectic transition in systems of soft parallel spherocylinders. Phys Rev A. Nov 15 1992;46(10):6541–6549. doi:10.1103/physreva.46.6541

85. Valdés-Tresanco MS, Valdés-Tresanco ME, Valiente PA, Moreno E. gmx_MMPBSA: A New Tool to Perform End-State Free Energy Calculations with GROMACS. J Chem Theory Comput. Oct 12 2021;17(10):6281–6291. doi:10.1021/acs.jctc.1c00645

