## Extended Data Figures-1-10 for "(2R,6R)-Hydroxynorketamine elicits rapid antidepressant effects by promoting astrocytic µ-δ opioid receptor heterodimerization"

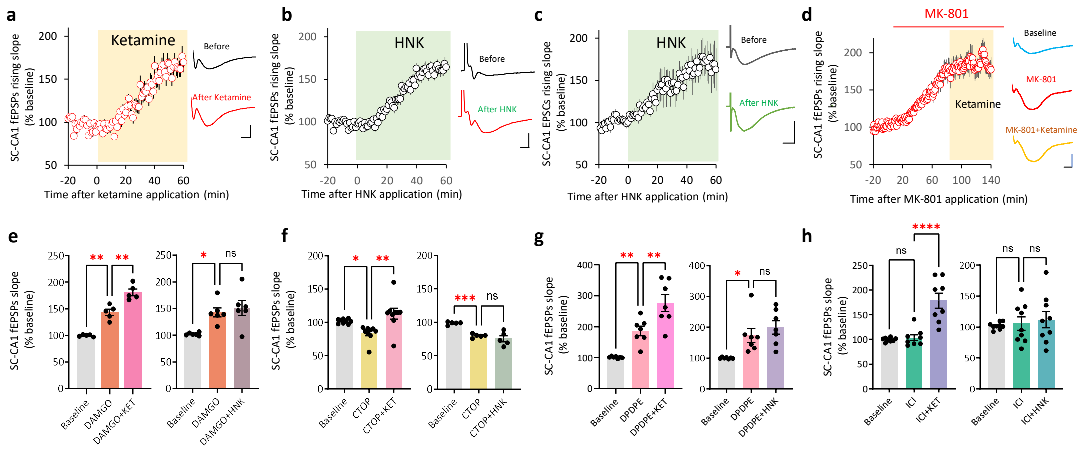


**Extended Data Fig. 1. Both ketamine and HNK potentiate Schaffer collateral – CA1 field EPSPs or EPSCs, with HNK-mediated synaptic potentiation dependent on opioid receptor activation**

**a,** The effect of ketamine on SC-CA1 fEPSPs, recorded in the stratum radiatum of CA1 in acutely prepared mouse hippocampal slices. Ketamine (10 μM) was applied to the bath. Left, data are presented as the time course of SC-CA1 fEPSPs slope before and after ketamine application (yellow shading) in artificial cerebrospinal fluid (ACSF). Ketamine increased SC-CA1 fEPSPs slope to 162.1 ± 2.1% of baseline (the comparison was measured by last 5 min of drug application with last 5 min of baseline, here and in all other electrophysiological experiments in the study).

**b,** HNK bath application significantly increased SC-CA1 fEPSPs slope by 161.9 ± 5.9% of baseline.

**c,** Whole-cell patch clamp experiments show that HNK increased SC-CA1 EPSCs slope by 163.2 ± 16.2% of baseline

**d,** MK-801 (10 μM) increased SC-CA1 fEPSPs slope and occluded the effect of ketamine (MK-801:167.8 ± 17.7% of baseline; ketamine plus MK-801:184.8 ± 20.5% of baseline).

**e**–**h**, data summaries of experiments showed in Fig. 1a–d.

Scale bar: 5 ms, 0.2 mV in a–c; 75 ms,100 pA in d.

Data are presented as mean ± SEM. *$P < 0.05$, **$P < 0.01$, ***$P < 0.001$, ****$P < 0.0001$. ns, not significant (statistical analyses and n numbers see Supplementary Table 1).

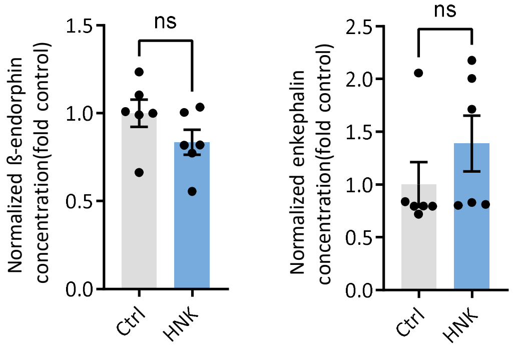


**Extended Data Fig. 2. HNK does not elevate β-endorphin and enkephalin level in the hippocampus**

No significant alterations in hippocampal β-endorphin and enkephalin levels were observed at 1 h following intraperitoneal injection of HNK (10 mg·kg$^{-1}$). Each dot in the bar graphs represents an individual mouse.

Data are presented as mean ± SEM. ns, not significant (statistical analyses and n numbers see Supplementary Table 1).

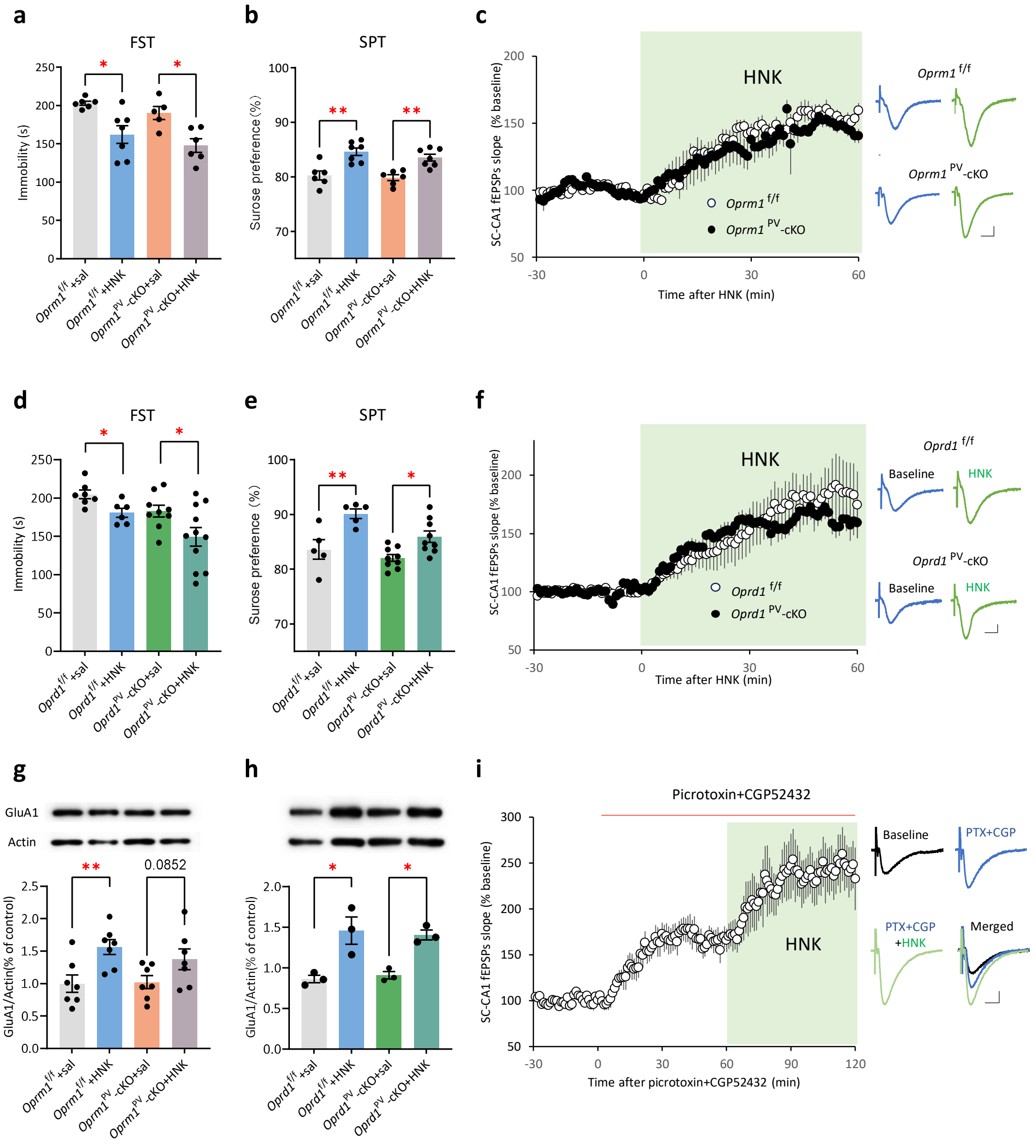


**Extended Data Fig. 3. Deletion of MOR or DOR in parvalbumin-positive (PV) neurons does not attenuate the antidepressant-like actions of HNK**

**a–b,** HNK (10 mg·kg$^{-1}$, i.p.) decreased immobility time in FST(a) and increased sucrose preference in SPT (b) in both *Oprm1*$^{PV}$-cKO mice and *Oprm1*$^{flox/flox}$ controls. **c,** HNK potentiated SC-CA1 fEPSPs in hippocampal slices from *Oprm1*$^{PV}$-cKO and *Oprm1*$^{flox/flox}$ mice. Left: time course of fEPSPs; Right, representative traces.

**d–e,** HNK (10 mg·kg$^{-1}$, i.p.) decreased immobility time in FST(d) and increased sucrose preference in SPT (e) in *Oprd1*$^{PV}$-cKO and *Oprd1*$^{flox/flox}$ mice.

**f,** HNK (10 μM) potentiated SC-CA1 fEPSPs in hippocampal slices from *Oprd1*$^{PV}$-cKO and *Oprd1*$^{flox/flox}$ mice. Left: time course of fEPSPs; Right, representative traces.

**g,** HNK (10 μM) increased GluA1 abundance in hippocampal slices from *Oprm1*$^{PV}$-cKO and *Oprm1*$^{flox/flox}$ mice.

**h,** HNK (10 μM) increased GluA1 abundance in hippocampal slices from *Oprd1*$^{PV}$-cKO and *Oprd1*$^{flox/flox}$ mice.

**i,** Blockade of $GABA_A$ and $GABA_B$ receptors by picrotoxin (100 μM) and CGP52432 (4 μM) enhanced SC-CA1 fEPSPs, and HNK induced further potentiation of SC-CA1 fEPSPs in the presence of picrotoxin and CGP. Left: time course of fEPSPs; Right, representative traces.

Scale bars in c, f, and i: 5 ms, 0.1 mV. Each dot in the bar graphs represents an individual mouse in (a, b, d, e), or independent Western blot experiment in (g, h).

Data are presented as mean ± SEM. *$P < 0.05$, **$P < 0.01$, ***$P < 0.001$, ****$P < 0.0001$. ns, not significant (statistical analyses and n numbers see Supplementary Table 1).

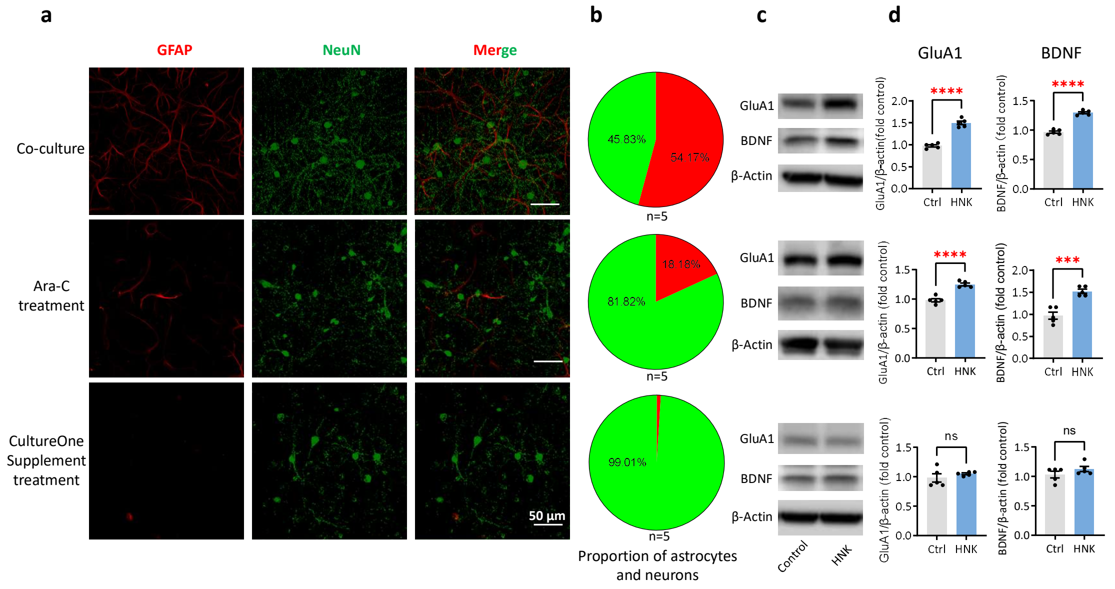


**Extended Data Fig. 4. Astrocytes are required for HNK-induced upregulation of BDNF and GluA1**

**a,** Immunofluorescence staining of GFAP (marker of astrocyte) and NeuN (marker of neuron) in hippocampal cell cultures with standard medium (upper), AraC (to suppress glia, middle), or CultureOne (glia-free, lower).

**b,** Pie charts showing neuron/astrocyte proportions in the three culture conditions.

**c–d,** Representative Western blots (c) and quantification (d) of HNK's effects on GluA1 and BDNF in the three culture conditions. Each dot in the bar graph represents an independent experiment.

Scale bars in a: 50 μm, Data are presented as mean ± SEM. *$P < 0.05$, **$P < 0.01$, ***$P < 0.001$, ****$P < 0.0001$. ns, not significant (statistical analyses and n numbers see Supplementary Table 1).

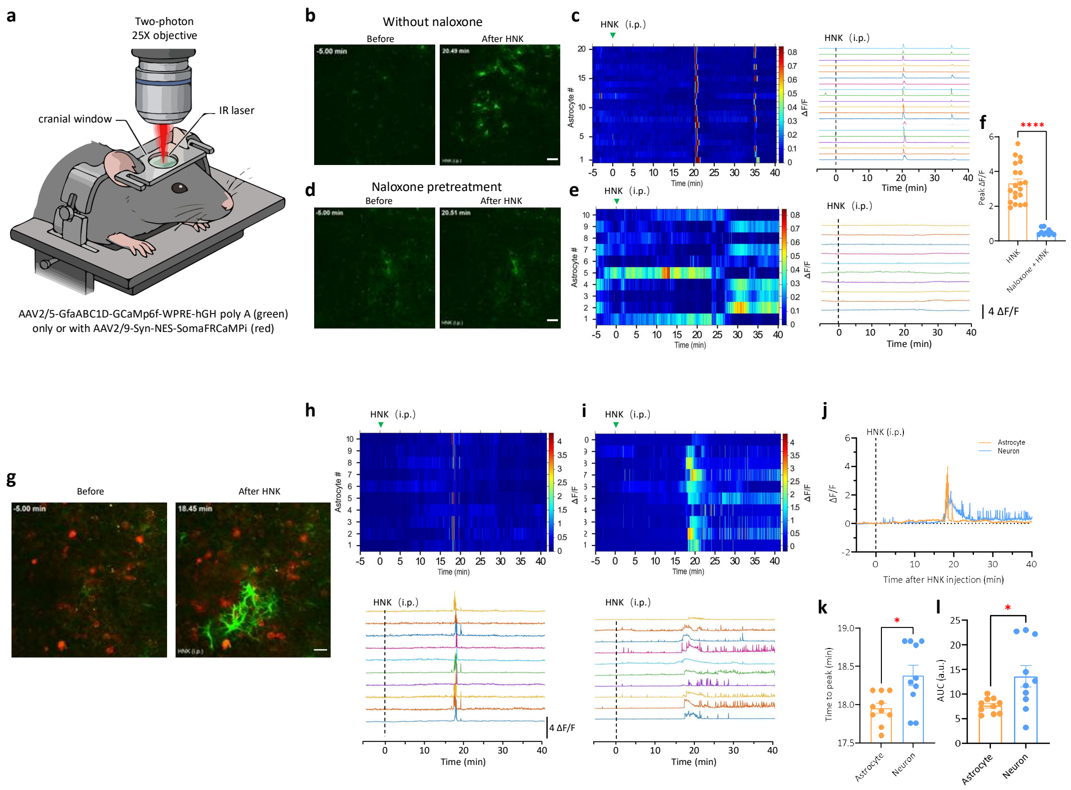


**Extended Data Fig. 5. HNK induces robust calcium transients in hippocampal CA1 astrocytes *in vivo*. This glial calcium response peaks significantly earlier than HNK-elicited calcium spikes in adjacent neurons and is fully abolished by naloxone pretreatment.**

**a,** Schematic diagram illustrating the setup for *in vivo* dual-color two-photon imaging of hippocampal astrocytic and neuronal calcium activity in lightly anesthetized mice. Schematic overview of the viral constructs used for astrocyte-specific GCaMP6f expression and neuron-specific expression of soma-restricted red calcium indicator (RCaMPi).

**b,** Representative fluorescence micrographs of GCaMP6f signals in the hippocampal CA1 region from a sample mouse, showing robust calcium elevation following intraperitoneal injection of 20 mg·kg$^{-1}$ HNK. Scale bar: 20 µm.

**c,** Time-resolved heatmap (left) and traces (right) of ΔF/F values for individual CA1 astrocytes (shown in b) in response to a single dose of HNK.

**d,** Representative GCaMP6f fluorescence micrographs showing hippocampal astrocytic calcium signals before and after HNK injection in mice pretreated with naloxone (1 mg·kg$^{-1}$, i.p., 20 min prior to HNK administration).

**e,** In mice pretreated with naloxone (1 mg·kg$^{-1}$, i.p., 20 min prior to HNK administration), Time-resolved heatmap (left) and traces (right) of ΔF/F values for individual CA1 astrocytes (shown in b) in response to a single dose of HNK.

**f,** Quantitative analysis of astrocytic calcium ΔF/F responses to HNK stimulation with or without naloxone pretreatment.

**g,** Representative dual-color two-photon images showing dynamic calcium responses of GCaMP6f-labeled astrocytes (green) and soma-restricted RCaMP-labeled neurons (red) in the hippocampal CA1 region following intraperitoneal injection of 20 mg·kg$^{-1}$ HNK. Scale bar: 20 μm.

**h,** Time-resolved heatmap (upper) and individual fluorescence traces (lower) of ΔF/F values for HNK-evoked calcium activity in individual astrocytes (green channel, shown in i).

**i,** Time-resolved heatmap (upper) and individual fluorescence traces (lower) of ΔF/F values for HNK-evoked calcium activity in individual neurons (red channel, shown in i).

**j,** Averaged ΔF/F traces illustrating temporal calcium activity profiles of HNK-stimulated astrocytes (n = 10, yellow) and neurons (n = 10, blue).

**k,** Quantitative comparison of the time-to-peak of ΔF/F signals in HNK-activated astrocytes and neurons. HNK-elicited astrocytic calcium transients peaked significantly earlier than neuronal calcium responses.

**l,** Quantitative analysis of the area under the curve (AUC) for calcium transients in HNK-stimulated astrocytes and neurons.

Data are presented as mean ± SEM. $*P < 0.05$, $**P < 0.01$, $***P < 0.001$, $****P < 0.0001$. ns, not significant (statistical analyses and n values are provided in Supplementary Table 1)

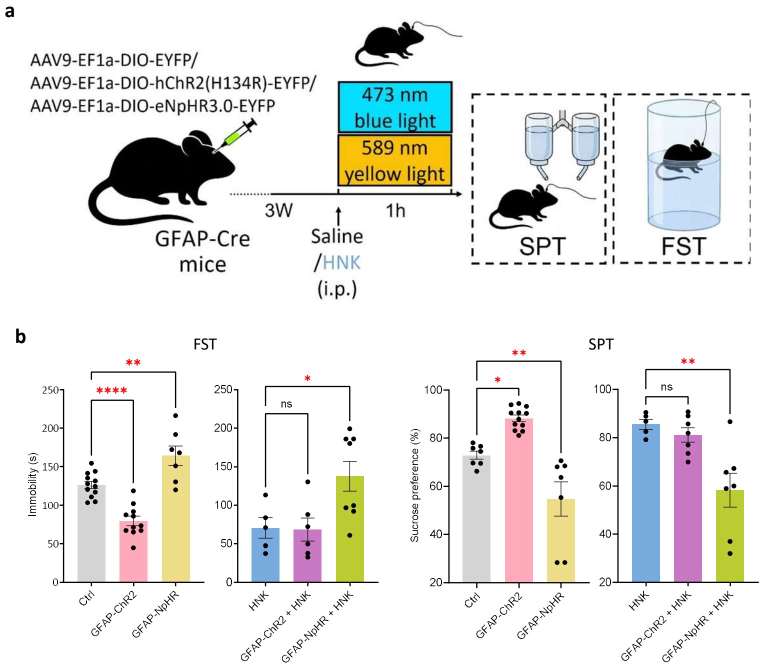


**Extended Data Fig. 6. Optogenetic activation hippocampal astrocytes occludes the antidepressant-like effects of HNK, whereas optogenetic inhibition of hippocampal astrocytes abolishes it.**

**a,** Experimental design of optogenetic manipulation.

**b,** Optogenetic activation (by 473 nm blue light) of hippocampal astrocytes (transfected with AAV9-EF1a-DIO-hChR2(H134R)-EYFP) mimicked and occluded the antidepressant-like effects of HNK in GFAP-Cre mice, whereas optogenetic inhibition (with 589 nm yellow light) of hippocampal astrocytes (transfected with AAV9-EF1a-DIO-eNpHR3.0-EYFP) induced depression-like effects in FST and SPT and blocked the antidepressant-like effects of HNK. Control GFAP-Cre mice were transfected with AAV9-EF1a-DIO-EYFP.

Data are presented as mean ± SEM. $*P < 0.05$, $**P < 0.01$, $***P < 0.001$, $****P < 0.0001$. ns, not significant (statistical analyses and n numbers see Supplementary Table 1).

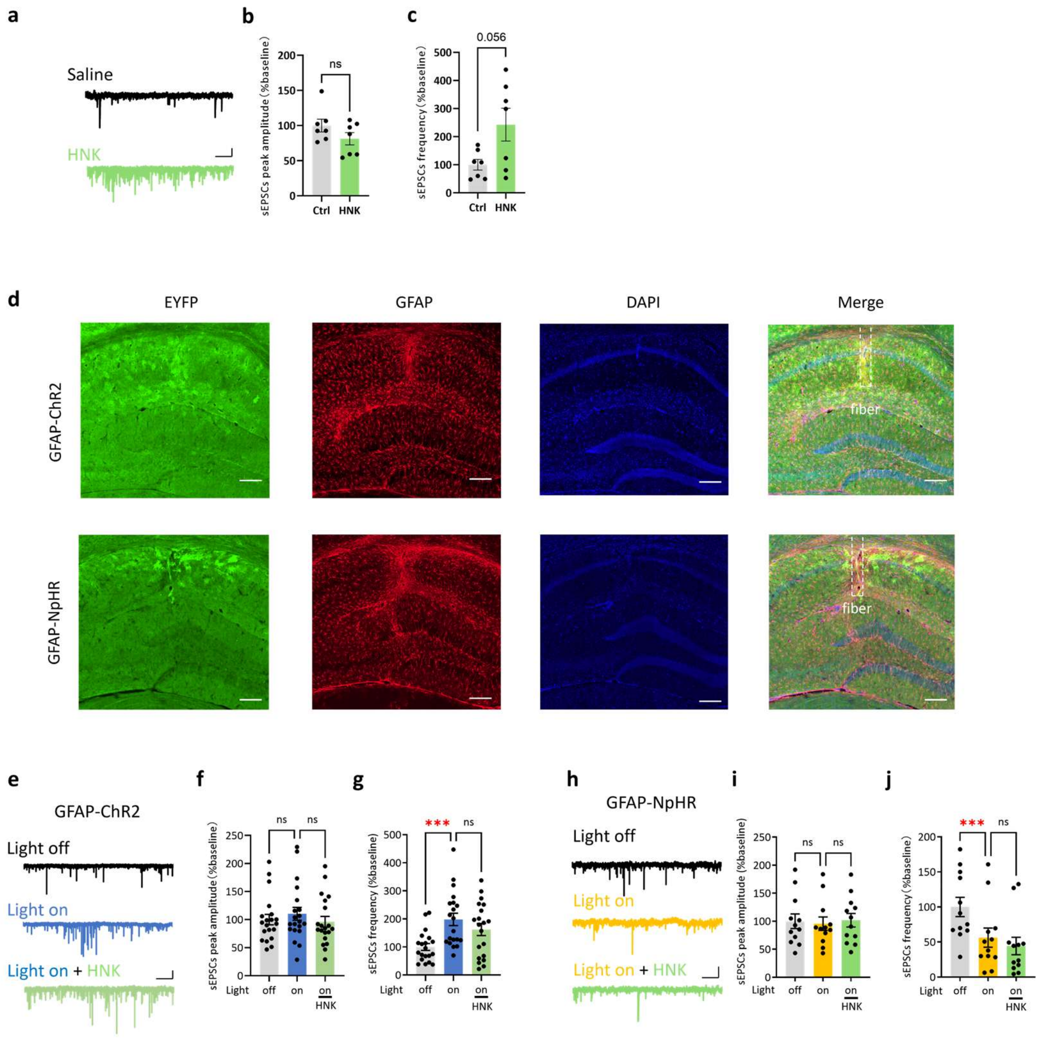


**Extended Data Fig. 7. Optogenetic activation hippocampal astrocytes occludes the effect of HNK in potentiation of excitatory synapses, whereas optogenetic inhibition of hippocampal astrocytes abolishes it**

**a–c,** Representative traces (a) and data summary (b, c) of sEPSCs recorded at CA1 pyramidal cells in hippocampal slices from wild-type mice before and after HNK (10 μM) perfusion, showing that HNK increased the frequency but not amplitude of sEPSCs from CA1 cells.

**d,** Immunofluorescence staining of EYFP, GFAP in hippocampal slices from GFAP-Cre mice infected with AAV9-EF1a-DIO-hChR2(H134R)-EYFP or AAV9-EF1a-DIO-eNpHR3.0-EYFP.

**e–g,** Representative traces (e) and data summary (f, g) of sEPSCs recorded at CA1 cells in hippocampal slices from GFAP-Cre mice infected with AAV9-EF1a-DIO-hChR2(H134R)-EYFP before, after turning blue light (473 nm) on, and after HNK application on top of blue light, showing that photo-activation of hippocampal astrocytes increased the frequency but not peak amplitude of sEPSCs from CA1 cells, and occluded HNK-induced increase in sEPSCs frequency.

**h–j,** Representative traces (h) and data summary (i, j) of sEPSCs recorded from CA1 cells in hippocampal slices from GFAP-Cre mice infected with AAV9-EF1a-DIO-eNpHR3.0-EYFP before, during yellow light (589 nm) illumination, and following subsequent HNK application under continued light exposure. These results demonstrate that photo-inhibition of hippocampal astrocytes did not alter the peak amplitude but reduced the frequency of sEPSCs recorded from CA1 cells, and abolished HNK-induced increase in sEPSCs frequency.

Scale bars: 100 μm in d; 2 s, 10 pA in a, e, h.

Data are presented as mean ± SEM. *$P < 0.05$, **$P < 0.01$, ***$P < 0.001$, ****$P < 0.0001$. ns, not significant (statistical analyses and n numbers see Supplementary Table 1).

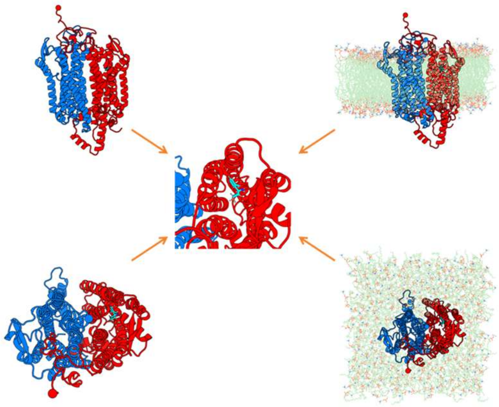

**Extended Data Fig. 8**. **Transmembrane structure of the MOR-DOR heterodimers and its interaction with HNK used in the simulations**

The atomistic structure of the MOR-DOR heterodimers is shown in side view (top) and top view (bottom), with the corresponding transmembrane model displayed on the right. The binding pocket of HNK is highlighted at the center. The initial heterodimers structure and ketamine binding site were predicted using AlphaFold3. For clarity, the MOR is colored red, and the DOR is colored blue.

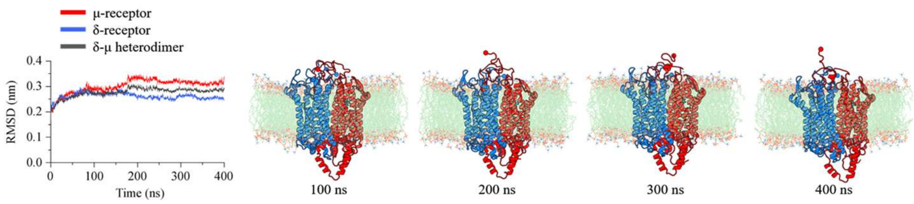


**Extended Data Fig. 9. Optimization of the MOR-DOR heterodimers with HNK**

Time evolution of the root mean square deviation (RMSD) for the MOR-DOR heterodimers, as well as for the individual MOR and DOR, relative to their initial states shown in Extended Data Fig.8. Representative snapshots from the simulations are displayed at 100 ns intervals. For clarity, the MOR and DOR are colored in red and blue, respectively.

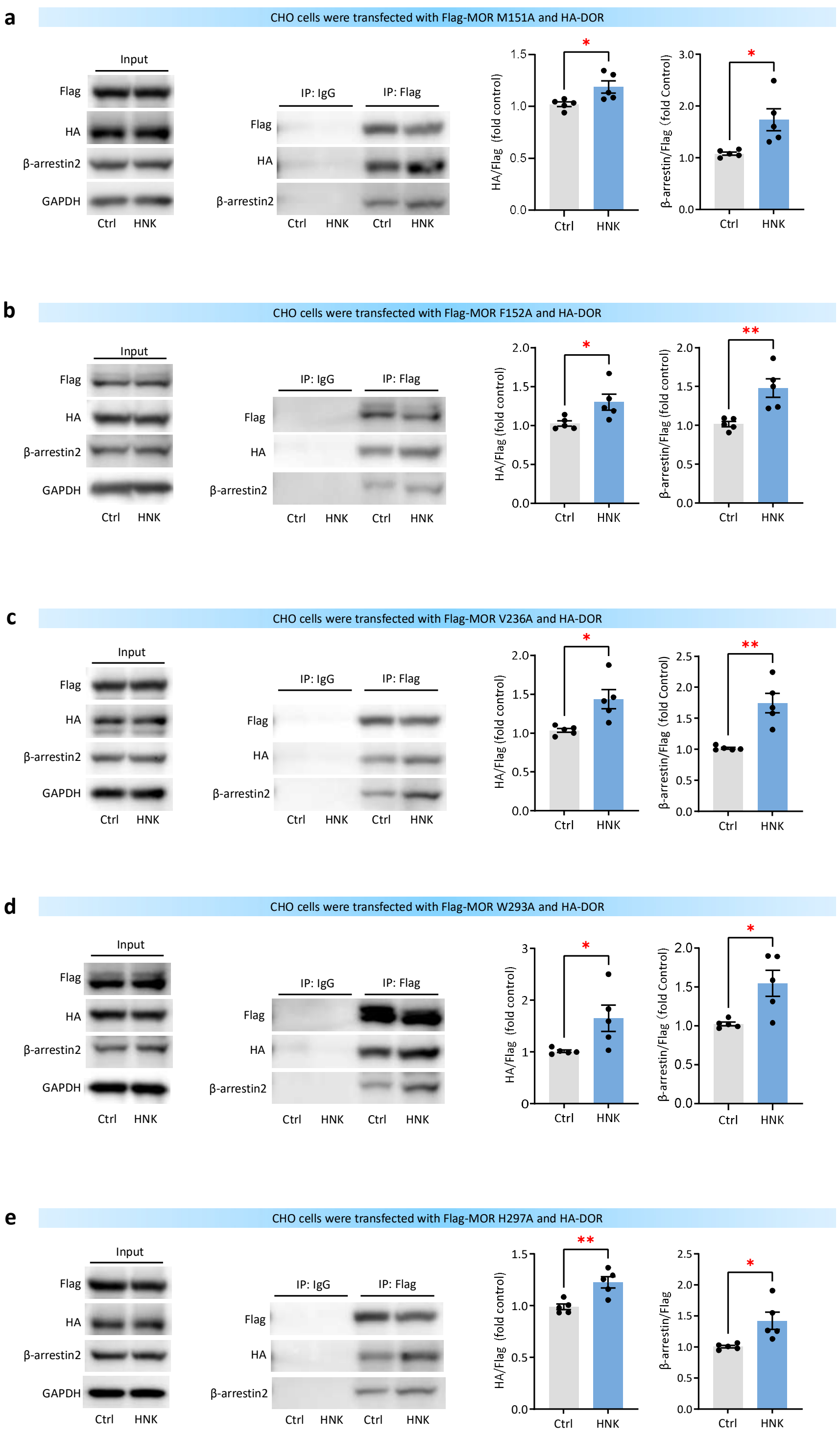


**Extended Data Fig. 10. M151A, F152A, V236A, W293A, or H297A point mutations of the MOR do not abolish the ability of HNK to promote MOR-DOR heterodimers formation and β-arrestin2 recruitment.**

CHO cells were transfected with Flag-tagged mutant MOR and HA-tagged wild-type

DOR. For each panel: Left: The input lysate of the CHO cells treated with HNK or vehicle (Ctrl) for 2 h were immunoblotted with antibodies against Flag, HA, β-arrestin2, or GAPDH; Middle: Lysates were immunoprecipitated with antibodies against IgG (negative control) or Flag, followed by immunoblotting with antibodies against Flag, HA, or β-arrestin2; Right: Quantification summary of the middle panel, showing the levels of HA-DOR or β-arrestin2 co-immunoprecipitated with Flag-MOR.

**a,** Transfection with Flag-MOR (M151A) and HA-DOR.

**b,** Transfection with Flag-MOR (F152A) and HA-DOR.

**c,** Transfection with Flag-MOR (V236A) and HA-DOR.

**d,** Transfection with Flag-MOR (W293A) and HA-DOR.

**e,** Transfection with Flag-MOR (H297A) and HA-DOR.

Data are presented as mean ± SEM. $*P < 0.05$, $**P < 0.01$, $***P < 0.001$, $****P < 0.0001$. ns, not significant (statistical analyses and n numbers see Supplementary Table 1).
