## Extended Data tables and movies for "(2R,6R)-Hydroxynorketamine elicits rapid antidepressant effects by promoting astrocytic µ-δ opioid receptor heterodimerization"

**Table S1.**

| Figure | Sample | Sample size | Statistic method | Statistic results |
|---|---|---|---|---|
| 1b | (SC-CA1 fEPSPs slope) baseline vs ketamine (10 μM) | n = 7 slices (5 mice) | Paired t test | *P* < 0.0001 |
| | (SC-CA1 fEPSPs slope) ketamine (10 μM) vs ketamine+HNK (10 μM) | n = 7 slices (5 mice) | Paired t test | *P* < 0.0001 |
| 1c | (SC-CA1 fEPSPs slope) baseline vs ketamine (10 μM) | n = 4 slices (4 mice) | Paired t test | *P* < 0.0001 |
| | (SC-CA1 fEPSPs slope) ketamine (10 μM) vs ketamine (30 μM) | n = 4 slices (4 mice) | Paired t test | *P* = 0.13 |
| 1d | (SC-CA1 fEPSPs slope) baseline vs MK-801 (10 μM) | n = 5 slices (4 mice) | Paired t test | *P* < 0.0001 |
| | (SC-CA1 fEPSPs slope) MK-801 (10 μM) vs MK-801+HNK (10 μM) | n = 5 slices (4 mice) | Paired t test | *P* < 0.0001 |
| 1e and S1e | (SC-CA1 fEPSPs slope) baseline vs DAMGO (1 μM, ketamine group) | n = 5 slices (4 mice) | Ordinary one-way ANOVA | *P* = 0.0023 |
| | (SC-CA1 fEPSPs slope) DAMGO (1 μM) vs DAMGO+ketamine (10 μM) | n = 5 slices (4 mice) | Ordinary one-way ANOVA | *P* = 0.0074 |
| | (SC-CA1 fEPSPs slope) baseline vs DAMGO (1 μM, HNK group) | n = 6 slices (5 mice) | Ordinary one-way ANOVA | *P* = 0.0162 |
| | (SC-CA1 fEPSPs slope) DAMGO (1 μM) vs DAMGO+HNK (10 μM) | n = 6 slices (5 mice) | Ordinary one-way ANOVA | *P* = 0.8171 |
| 1f and S1f | (SC-CA1 fEPSPs slope) baseline vs CTOP (1 μM, ketamine group) | n = 9 slices (7 mice) | Ordinary one-way ANOVA | *P* = 0.0441 |
| | (SC-CA1 fEPSPs slope) CTOP (1 μM) vs CTOP+ketamine (10 μM) | n = 9 slices (7 mice) | Ordinary one-way ANOVA | *P* = 0.0012 |
| | (SC-CA1 fEPSPs slope) baseline vs CTOP (1 μM, HNK group) | n = 5 slices (4 mice) | Ordinary one-way ANOVA | *P* = 0.0010 |
| | (SC-CA1 fEPSPs slope) CTOP (1 μM) vs CTOP+HNK (10 μM) | n = 5 slices (4 mice) | Ordinary one-way ANOVA | *P* = 0.4750 |
| 1g and S1g | (SC-CA1 fEPSPs slope) baseline vs DPDPE (1 μM, ketamine group) | n = 7 slices (6 mice) | Ordinary one-way ANOVA | *P* = 0.0070 |
| | (SC-CA1 fEPSPs slope) DPDPE (1 μM) vs DPDPE+ketamine (10 μM) | n = 7 slices (6 mice) | Ordinary one-way ANOVA | *P* = 0.0043 |
| | (SC-CA1 fEPSPs slope) baseline vs DPDPE (1 μM, HNK group) | n = 7 slices (4 mice) | Ordinary one-way ANOVA | *P* = 0.0374 |
| | (SC-CA1 fEPSPs slope) DPDPE (1 μM) vs DPDPE+HNK (10 μM) | n = 7 slices (4 mice) | Ordinary one-way ANOVA | *P* = 0.3894 |
| 1h and S1h | (SC-CA1 fEPSPs slope) baseline vs ICI (1 μM, ketamine group) | n = 8 slices (7 mice) | Ordinary one-way ANOVA | *P* = 0.9415 |
| | (SC-CA1 fEPSPs slope) ICI (1 μM) vs ICI+ketamine (10 | n = 8 slices (7 mice) | Ordinary one-way ANOVA | *P* < 0.0001 |

| | | | | |
|---|---|---|---|---|
| | μM) | | | |
| | (SC-CA1 fEPSPs slope)baseline vs ICI (1 μM, HNK group) | n = 9 slices (6 mice) | Ordinary one-way ANOVA | $P = 0.9616$ |
| | (SC-CA1 fEPSPs slope) ICI (1 μM) vs ICI+HNK (10 μM) | n = 9 slices (6 mice) | Ordinary one-way ANOVA | $P = 0.8764$ |
| 1j | (FST) saline vs HNK | n = 6, 6 | Ordinary one-way ANOVA | $P = 0.0134$ |
| | (FST) saline vs DAMGO | n = 6, 6 | Ordinary one-way ANOVA | $P = 0.0184$ |
| | (FST) HNK vs CTOP+HNK | n = 6, 6 | Ordinary one-way ANOVA | $P = 0.0417$ |
| 1k | (SPT) saline vs HNK | n = 6, 6 | Ordinary one-way ANOVA | $P = 0.0428$ |
| | (SPT) saline vs DAMGO | n = 6, 6 | Ordinary one-way ANOVA | $P = 0.0270$ |
| | (SPT) HNK vs CTOP+HNK | n = 6, 6 | Ordinary one-way ANOVA | $P < 0.0001$ |
| 1l | (One stranger) Ctrl vs HNK | n = 7, 7 | Ordinary one-way ANOVA | $P = 0.0003$ |
| | (One stranger) Ctrl vs DAMGO | n = 7, 6 | Ordinary one-way ANOVA | $P = 0.0011$ |
| | (One stranger) HNK vs CTOP+HNK | n = 7, 6 | Ordinary one-way ANOVA | $P = 0.0024$ |
| | (Two strangers) Ctrl vs HNK | n = 6, 6 | Ordinary one-way ANOVA | $P = 0.0026$ |
| | (Two strangers) Ctrl vs DAMGO | n = 6, 6 | Ordinary one-way ANOVA | $P = 0.0014$ |
| | (Two strangers) HNK vs CTOP+HNK | n = 6, 6 | Ordinary one-way ANOVA | $P = 0.0385$ |
| 1m | (FST) Ctrl vs MOR KO | n = 6, 6 | Unpaired t test | $P = 0.0062$ |
| 1n | (SPT) Ctrl vs MOR KO | n = 8, 8 | Unpaired t test | $P = 0.0497$ |
| 1o | (FST) MOR KO vs MOR KO+HNK | n = 5, 5 | Unpaired t test | $P = 0.3429$ |
| 1p | (SPT) MOR KO vs MOR KO+HNK | n = 8, 8 | Unpaired t test | $P = 0.5863$ |
| 1q | (FST) Ctrl vs HNK | n = 6, 6 | Ordinary one-way ANOVA | $P = 0.0045$ |
| | (FST) Ctrl vs DPDPE | n = 6, 8 | Ordinary one-way ANOVA | $P = 0.0019$ |
| | (FST) HNK vs ICI+HNK | n = 6, 8 | Ordinary one-way ANOVA | $P = 0.0006$ |
| 1r | (SPT) Ctrl vs HNK | n = 8, 8 | Ordinary one-way ANOVA | $P = 0.0301$ |
| | (SPT) Ctrl vs DPDPE | n = 8, 8 | Ordinary one-way ANOVA | $P = 0.0009$ |
| | (SPT) HNK vs ICI+HNK | n = 8, 8 | Ordinary one-way ANOVA | $P = 0.0004$ |
| 1s | (GluA1/Actin) Ctrl vs HNK | n = 6, 6 | Ordinary one-way ANOVA | $P = 0.0017$ |
| | (GluA1/Actin) Ctrl vs DAMGO | n = 6, 6 | Ordinary one-way ANOVA | $P < 0.0001$ |
| | (GluA1/Actin) DAMGO vs DAMGO+HNK | n = 6, 6 | Ordinary one-way ANOVA | $P = 0.3905$ |
| 1t | (GluA1/Actin) Ctrl vs HNK | n = 6, 6 | Ordinary one-way ANOVA | $P = 0.0003$ |
| | (GluA1/Actin) HNK vs CTOP+HNK | n = 6, 6 | Ordinary one-way ANOVA | $P = 0.0011$ |
| | (GluA1/Actin) CTOP vs CTOP+HNK | n = 6, 6 | Ordinary one-way ANOVA | $P = 0.9901$ |

| | | | | |
|---|---|---|---|---|
| 1u | (GluA1/Actin) Ctrl vs DPDPE | n = 7, 7 | Ordinary one-way ANOVA | *P* = 0.0051 |
| | (GluA1/Actin) Ctrl vs HNK | n = 7, 7 | Ordinary one-way ANOVA | *P* = 0.0137 |
| | (GluA1/Actin) DPDPE vs HNK | n = 7, 7 | Ordinary one-way ANOVA | *P* = 0.9753 |
| | (GluA1/Actin) HNK vs DPDPE+HNK | n = 7, 7 | Ordinary one-way ANOVA | *P* = 0.8980 |
| 1v | (GluA1/Actin) Ctrl vs ICI | n = 3, 3 | Ordinary one-way ANOVA | *P* = 0.1824 |
| | (GluA1/Actin) Ctrl vs HNK | n = 3, 3 | Ordinary one-way ANOVA | *P* = 0.0239 |
| | (GluA1/Actin) ICI vs ICI+HNK | n = 3, 3 | Ordinary one-way ANOVA | *P* = 0.9978 |
| | (GluA1/Actin) HNK vs ICI+HNK | n = 3, 3 | Ordinary one-way ANOVA | *P* = 0.0018 |
| 2b | (FST) WT+saline vs CORT+saline | n = 8, 7 | Ordinary one-way ANOVA | *P* = 0.0285 |
| | (FST) CORT+saline vs CORT+HNK | n = 7, 5 | Ordinary one-way ANOVA | *P* = 0.0156 |
| | (FST) CORT+HNK vs CORT+CYM | n = 5, 7 | Ordinary one-way ANOVA | *P* = 0.9993 |
| | (FST) CORT+CYM vs CORT+CYM+HNK | n = 7, 5 | Ordinary one-way ANOVA | *P* > 0.9999 |
| | (FST) CORT+PEP vs CORT+PEP+HNK | n = 5, 7 | Ordinary one-way ANOVA | *P* > 0.9999 |
| 2c | (TST) WT+saline vs CORT+saline | n = 13, 14 | Ordinary one-way ANOVA | *P* < 0.0001 |
| | (TST) CORT+saline vs CORT+HNK | n = 14, 15 | Ordinary one-way ANOVA | *P* = 0.0436 |
| | (TST) CORT+saline vs CORT+CYM | n = 14, 12 | Ordinary one-way ANOVA | *P* = 0.0065 |
| | (TST) CORT+CYM vs CORT+CYM+HNK | n = 12, 7 | Ordinary one-way ANOVA | *P* = 0.9982 |
| | (TST) CORT+PEP vs CORT+PEP+HNK | n = 10, 10 | Ordinary one-way ANOVA | *P* > 0.9999 |
| 2d | (One stranger) WT+saline vs CORT+saline | n = 12, 11 | Ordinary one-way ANOVA | *P* < 0.0001 |
| | (One stranger) CORT+saline vs CORT+HNK | n = 11, 14 | Ordinary one-way ANOVA | *P* = 0.0010 |
| | (One stranger) CORT+HNK vs CORT+CYM | n = 14, 6 | Ordinary one-way ANOVA | *P* = 0.9376 |
| | (One stranger) CORT+CYM vs CORT+CYM+HNK | n = 6, 6 | Ordinary one-way ANOVA | *P* > 0.9999 |
| | (One stranger) CORT+PEP vs CORT+PEP+HNK | n = 6, 6 | Ordinary one-way ANOVA | *P* = 0.6833 |
| | (Two strangers) WT+saline vs CORT+saline | n = 14, 6 | Ordinary one-way ANOVA | *P* < 0.0001 |
| | (Two strangers) CORT+saline vs CORT+HNK | n = 6, 6 | Ordinary one-way ANOVA | *P* = 0.0007 |
| | (Two strangers) CORT+saline vs CORT+CYM | n = 6, 6 | Ordinary one-way ANOVA | *P* = 0.0447 |
| | (Two strangers) | n = 6, 6 | Ordinary one- | *P* = 0.9643 |

| | | | | |
|---|---|---|---|---|
| | CORT+CYM vs CORT+CYM+HNK | | way ANOVA | |
| | (Two strangers) CORT+PEP vs CORT+PEP+HNK | n = 6, 6 | Ordinary one-way ANOVA | *P* = 0.9558 |
| 2e | (SC-CA1 fEPSPs slope) baseline vs CYM (1 μM) | n = 13 slices (8 mice) | Paired t test | *P* = 0.0349 |
| | (SC-CA1 fEPSPs slope) CYM (1 μM) vs CYM+HNK (10 μM) | n = 13 slices (8 mice) | Paired t test | *P* = 0.6322 |
| 2f | (SC-CA1 fEPSPs slope) baseline vs PEP (50 nM) | n = 14 slices (8 mice) | Paired t test | *P* = 0.1783 |
| | (SC-CA1 fEPSPs slope) PEP (50 nM) vs PEP+HNK (10 μM) | n = 14 slices (8 mice) | Paired t test | *P* = 0.9795 |
| 2g | (GluA1/Actin) Ctrl vs HNK | n = 7, 7 | Ordinary one-way ANOVA | *P* = 0.0451 |
| | (GluA1/Actin) Ctrl vs CYM | n = 7, 5 | Ordinary one-way ANOVA | *P* = 0.0006 |
| | (GluA1/Actin) HNK vs CYM | n = 7, 5 | Ordinary one-way ANOVA | *P* = 0.1555 |
| | (GluA1/Actin) CYM vs CYM+HNK | n = 5, 5 | Ordinary one-way ANOVA | *P* = 0.9999 |
| 2h | (GluA1/Actin) Ctrl vs HNK | n = 8, 8 | Ordinary one-way ANOVA | *P* = 0.0410 |
| | (GluA1/Actin) HNK vs PEP+HNK | n = 8, 5 | Ordinary one-way ANOVA | *P* = 0.0060 |
| | (GluA1/Actin) PEP vs PEP+HNK | n = 5, 5 | Ordinary one-way ANOVA | *P* = 0.8948 |
| 2j | (MOR-DOR/GAPDH) Ctrl vs CUMS | n = 10, 10 | Ordinary one-way ANOVA | *P* = 0.0017 |
| | (MOR-DOR/GAPDH) CUMS vs CUMS+HNK 1 h | n = 10, 10 | Ordinary one-way ANOVA | *P* = 0.0238 |
| | (MOR-DOR/GAPDH) CUMS vs CUMS+HNK 2 h | n = 10, 10 | Ordinary one-way ANOVA | *P* = 0.0008 |
| | (MOR-DOR/GAPDH) CUMS vs CUMS+HNK 24 h | n = 10, 10 | Ordinary one-way ANOVA | *P* = 0.2399 |
| 2k | (MOR/GAPDH) Ctrl vs CUMS | n = 10, 10 | Ordinary one-way ANOVA | *P* = 0.8562 |
| | (MOR/GAPDH) CUMS vs CUMS+HNK 1 h | n = 10, 10 | Ordinary one-way ANOVA | *P* = 0.9880 |
| | (MOR/GAPDH) CUMS vs CUMS+HNK 2 h | n = 10, 10 | Ordinary one-way ANOVA | *P* = 0.9708 |
| | (MOR/GAPDH) CUMS vs CUMS+HNK 24 h | n = 10, 10 | Ordinary one-way ANOVA | *P* = 0.8323 |
| 2l | (DOR/GAPDH) Ctrl vs CUMS | n = 10, 10 | Ordinary one-way ANOVA | *P* = 0.1706 |
| | (DOR/GAPDH) CUMS vs CUMS+HNK 1 h | n = 10, 10 | Ordinary one-way ANOVA | *P* = 0.9995 |
| | (DOR/GAPDH) CUMS vs CUMS+HNK 2 h | n = 10, 10 | Ordinary one-way ANOVA | *P* = 0.9942 |
| | (DOR/GAPDH) CUMS vs CUMS+HNK 24 h | n = 10, 10 | Ordinary one-way ANOVA | *P* = 0.9958 |
| 3c | (BDNF/Actin) Neuron saline vs Neuron HNK | n = 5, 5 | Unpaired t test | *P* = 0.7095 |
| | (BDNF/Actin) Astrocyte saline vs Astrocyte HNK | n = 5, 5 | Unpaired t test | *P* = 0.0011 |

| | | | | |
|---|---|---|---|---|
| | (BDNF/Actin) Co-culture saline vs Co-culture HNK | n = 5, 5 | Unpaired t test | *P* = 0.0004 |
| 3d | (GluA1/Actin) Neuron saline vs Neuron HNK | n = 5, 5 | Unpaired t test | *P* = 0.7988 |
| | (GluA1/Actin) Astrocyte saline vs Astrocyte HNK | n = 5, 5 | Unpaired t test | *P* = 0.7064 |
| | (GluA1/Actin) Co-culture saline vs Co-culture HNK | n = 5, 5 | Unpaired t test | *P* = 0.0018 |
| 3g | (GluA1 punctum number ) Without Astrocytes saline vs HNK | n = 6, 6 | Unpaired t test | *P* = 0.9385 |
| | (GluA1 punctum number) With Astrocytes saline vs HNK | n = 6, 6 | Unpaired t test | *P* = 0.0003 |
| | (GluA1 punctum size) Without Astrocytes saline vs HNK | n = 6, 6 | Unpaired t test | *P* = 0.5596 |
| | (GluA1 punctum size ) With Astrocytes saline vs HNK | n = 6, 6 | Unpaired t test | *P* < 0.0001 |
| 3i | (Number of colocalization puncta per astrocyte) Ctrl vs DXMS | n = 4, 4 | Ordinary one-way ANOVA | *P* = 0.0390 |
| | (Number of colocalization puncta per astrocyte) DXMS vs DXMS+HNK | n = 4, 4 | Ordinary one-way ANOVA | *P* = 0.0005 |
| 3j | (FST) *Oprm1*$^{flox/flox}$+saline vs *Oprm1*$^{flox/flox}$+HNK | n = 8, 8 | Ordinary one-way ANOVA | *P* = 0.0249 |
| | (FST) *Oprm1*$^{GFAP}$-cKO+saline vs *Oprm1*$^{GFAP}$-cKO+HNK | n = 6, 5 | Ordinary one-way ANOVA | *P* = 0.9978 |
| | (SPT) *Oprm1*$^{flox/flox}$+saline vs *Oprm1*$^{flox/flox}$+HNK | n = 8, 8 | Ordinary one-way ANOVA | *P* = 0.0199 |
| | (SPT) *Oprm1*$^{GFAP}$-cKO+saline vs *Oprm1*$^{GFAP}$-cKO+HNK | n = 5, 5 | Ordinary one-way ANOVA | *P* = 0.9938 |
| 3k | (SC-CA1 fEPSPs slope) *Oprm1*$^{flox/flox}$+HNK vs *Oprm1*$^{GFAP}$-cKO+HNK) | n = 8 slices (4 mice) | Unpaired t test | *P* < 0.0001 |
| 3l | (GluA1/Actin) *Oprm1*$^{flox/flox}$+saline vs *Oprm1*$^{flox/flox}$+HNK | n = 10, 10 | Unpaired t test | *P* = 0.0061 |
| | (GluA1/Actin) *Oprm1*$^{GFAP}$-cKO+saline vs *Oprm1*$^{GFAP}$-cKO+HNK | n = 8, 8 | Unpaired t test | *P* = 0.5171 |
| 3m | (FST) *Oprd1*$^{flox/flox}$+saline vs *Oprd1*$^{flox/flox}$+HNK | n = 7, 7 | Ordinary one-way ANOVA | *P* = 0.0007 |
| | (FST) *Oprd1*$^{GFAP}$-cKO+saline vs *Oprd1*$^{GFAP}$-cKO+HNK | n = 7, 7 | Ordinary one-way ANOVA | *P* = 0.8472 |
| | (SPT) *Oprd1*$^{flox/flox}$+saline vs *Oprd1*$^{flox/flox}$+HNK | n = 7, 7 | Ordinary one-way ANOVA | *P* = 0.0392 |
| | (SPT) *Oprd1*$^{flox/flox}$+saline vs *Oprd1*$^{GFAP}$-cKO+saline | n = 7, 7 | Ordinary one-way ANOVA | *P* = 0.8818 |

| | | | | |
|---|---|---|---|---|
| | (SPT) *Oprd1*$^{GFAP}$-cKO+saline vs *Oprd1*$^{GFAP}$-cKO+HNK | n = 7, 7 | Ordinary one-way ANOVA | *P* = 0.8252 |
| 3n | (SC-CA1 fEPSPs slope) *Oprd1*$^{flox/flox}$+HNK vs *Oprd1*$^{GFAP}$-cKO+HNK) | n = 8 slices (5 mice),<br>n = 9 slices (4 mice) | Unpaired t test | *P* < 0.0001 |
| 3o | (GluA1/Actin) *Oprd1*$^{flox/flox}$+saline vs *Oprd1*$^{flox/flox}$+HNK | n = 8, 8 | Ordinary one-way ANOVA | *P* = 0.0393 |
| | (GluA1/Actin) *Oprd1*$^{GFAP}$-cKO+saline vs *Oprd1*$^{GFAP}$-cKO+HNK | n = 8, 8 | Ordinary one-way ANOVA | *P* = 0.4869 |
| 4i | Dimer percentile (%) without Peptide (Ctrl vs HNK) | n = 18, 18 | Kruskal-Wallis test with Dunn's post hoc test | *P* < 0.0001 |
| | Dimer percentile (%) without Peptide (Ctrl vs CYM) | n = 18, 18 | Kruskal-Wallis test with Dunn's post hoc test | *P* < 0.0001 |
| | Dimer percentile (%) without Peptide (HNK vs CYM) | n = 18, 18 | Kruskal-Wallis test with Dunn's post hoc test | *P* = 0.3975 |
| | Dimer percentile (%) with Peptide (Ctrl vs CYM) | n = 18, 18 | Kruskal-Wallis test with Dunn's post hoc test | *P* > 0.9999 |
| | Dimer percentile (%) with Peptide (HNK vs CYM) | n = 18, 18 | Kruskal-Wallis test with Dunn's post hoc test | *P* > 0.9999 |
| 5g | HA/Flag Ctrl vs HNK | n = 5, 5 | Unpaired t test | *P* = 0.0354 |
| | β-arrestin/Flag Ctrl vs HNK | n = 5, 5 | Unpaired t test | *P* = 0.0003 |
| 5h | HA/Flag Ctrl vs HNK | n = 5, 5 | Unpaired t test | *P* = 0.9251 |
| | β-arrestin/Flag Ctrl vs HNK | n = 5, 5 | Unpaired t test | *P* = 0.2983 |
| 5i | HA/Flag Ctrl vs HNK | n = 5, 5 | Unpaired t test | *P* = 0.9361 |
| | β-arrestin/Flag Ctrl vs HNK | n = 5, 5 | Unpaired t test | *P* = 0.9626 |
| 5j | HA/Flag Ctrl vs HNK | n = 5, 5 | Unpaired t test | *P* = 0.5793 |
| | β-arrestin/Flag Ctrl vs HNK | n = 5, 5 | Unpaired t test | *P* = 0.7052 |
| 5l | (FST) WT (saline vs HNK) | n = 7, 7 | Paired t test | *P* = 0.0194 |
| | (FST) CUMS control AAV (saline vs HNK) | n = 8, 8 | Paired t test | *P* = 0.0011 |
| | (FST) CUMS+2mutations (saline vs HNK) | n = 7, 7 | Paired t test | *P* = 0.3957 |
| | (FST) CUMS+3mutations (saline vs HNK) | n = 8, 8 | Paired t test | *P* = 0.3119 |
| 5m | (SPT) WT (saline vs HNK) | n = 8, 8 | Paired t test | *P* = 0.0001 |
| | (SPT) CUMS control AAV (saline vs HNK) | n = 9, 9 | Paired t test | *P* < 0.0001 |
| | (SPT) CUMS+2mutations (saline vs HNK) | n = 9, 9 | Paired t test | *P* = 0.1160 |
| | (SPT) CUMS+3mutations (saline vs HNK) | n = 10, 10 | Paired t test | *P* = 0.9196 |
| S1a | (SC-CA1 fEPSPs slope) baseline vs ketamine (10 μM) | n = 7 slices (5 mice) | Paired t test | *P* < 0.0001 |
| S1b | (SC-CA1 fEPSPs slope) baseline vs HNK (10 μM) | n = 7 slices (5 mice) | Paired t test | *P* < 0.0001 |
| S1c | (SC-CA1 fEPSCs slope) baseline vs HNK (10 μM) | n = 8 slices (8 mice) | Paired t test | *P* < 0.0001 |

| | | | | |
|---|---|---|---|---|
| S1d | (SC-CA1 fEPSPs slope) baseline vs MK-801 (10 μM) | n = 20 slices (15 mice) | Paired t test | *P* < 0.0001 |
| | (SC-CA1 fEPSPs slope) MK-801 (10 μM) vs MK-801+ketamine (10 μM) | n = 20 slices (15 mice) | Ordinary one-way ANOVA | *P* = 0.8472 |
| S2 | (Normalized ß-endorphin Concentration) Ctrl vs HNK | n = 6, 6 | Unpaired t test | *P* = 0.1450 |
| | (Normalized enkephalin Concentration) Ctrl vs HNK | n = 6, 6 | Unpaired t test | *P* = 0.2766 |
| S3a | (FST) *Oprm1*$^{flox/flox}$+saline vs *Oprm1*$^{flox/flox}$+HNK | n = 6, 7 | Unpaired t test | *P* = 0.0080 |
| | (FST) *Oprm1*$^{PV}$-cKO+saline vs *Oprm1*$^{PV}$-cKO+HNK | n = 5, 6 | Unpaired t test | *P* = 0.0081 |
| S3b | (SPT) *Oprm1*$^{flox/flox}$+saline vs *Oprm1*$^{flox/flox}$+HNK | n = 6, 7 | Unpaired t test | *P* = 0.0022 |
| | (SPT) *Oprm1*$^{PV}$-cKO+saline vs *Oprm1*$^{PV}$-cKO+HNK | n = 6, 7 | Unpaired t test | *P* = 0.0012 |
| S3c | (SC-CA1 fEPSPs slope) *Oprm1*$^{flox/flox}$+HNK vs *Oprm1*$^{PV}$-cKO+HNK | n = 8 slices (4 mice), n = 6 slices (3 mice) | Unpaired t test | *P* = 0.3109 |
| S3d | (FST) *Oprd1*$^{flox/flox}$+saline vs *Oprd1*$^{flox/flox}$+HNK | n = 7, 6 | Unpaired t test | *P* = 0.0146 |
| | (FST) *Oprd1*$^{PV}$-cKO+saline vs *Oprd1*$^{PV}$-cKO+HNK | n = 9, 11 | Unpaired t test | *P* = 0.0406 |
| S3e | (SPT) *Oprd1*$^{flox/flox}$+saline vs *Oprd1*$^{flox/flox}$+HNK | n = 5, 5 | Unpaired t test | *P* = 0.0110 |
| | (SPT) *Oprd1*$^{PV}$-cKO+saline vs *Oprd1*$^{PV}$-cKO+HNK | n = 9, 9 | Unpaired t test | *P* = 0.0058 |
| S3f | (SC-CA1 fEPSPs slope) *Oprd1*$^{flox/flox}$+HNK vs *Oprd1*$^{PV}$-cKO+HNK | n = 5 slices (4 mice), n = 9 slices (5 mice) | Paired t test | *P* = 0.5194 |
| S3g | (GluA1/Actin) *Oprm1*$^{flox/flox}$+saline vs *Oprm1*$^{flox/flox}$+HNK | n = 7, 7 | Unpaired t test | *P* = 0.0081 |
| | (GluA1/Actin) *Oprm1*$^{PV}$-cKO+saline vs *Oprm1*$^{PV}$-cKO+HNK | n = 7, 7 | Unpaired t test | *P* = 0.0852 |
| S3h | (GluA1/Actin) *Oprd1*$^{flox/flox}$+saline vs *Oprd1*$^{flox/flox}$+HNK | n = 3, 3 | Unpaired t test | *P* = 0.0262 |
| | (GluA1/Actin) *Oprd1*$^{PV}$-cKO+saline vs *Oprd1*$^{PV}$-cKO+HNK | n = 3, 3 | Unpaired t test | *P* = 0.0028 |
| S3i | (SC-CA1 fEPSPs slope) baseline vs picrotoxin (100 μM)+CGP (4 μM) | n = 8 slices (6 mice) | Paired t test | *P* < 0.0001 |

| | | | | |
|---|---|---|---|---|
| | (SC-CA1 fEPSPs slope) picrotoxin (100 μM)+CGP (4 μM) vs picrotoxin (100 μM)+CGP (4 μM)+HNK | n = 8 slices (6 mice) | Paired t test | $P < 0.0001$ |
| S4d | (Co-culture GluA1/Actin) Ctrl vs HNK | n = 5, 5 | Unpaired t test | $P < 0.0001$ |
| | (Co-culture BDNF/Actin) Ctrl vs HNK | n = 5, 5 | Unpaired t test | $P < 0.0001$ |
| | (Ara-C treatment GluA1/Actin) Ctrl vs HNK) | n = 5, 5 | Unpaired t test | $P < 0.0001$ |
| | (Ara-C treatment BDNF/Actin) Ctrl vs HNK) | n = 5, 5 | Unpaired t test | $P = 0.0004$ |
| | (CultureOne Supplement Treatment GluA1/Actin) Ctrl vs HNK | n = 5, 5 | Unpaired t test | $P = 0.3541$ |
| | (CultureOne Supplement Treatment BDNF/Actin) Ctrl vs HNK | n = 5, 5 | Unpaired t test | $P = 0.2161$ |
| S5f | (Peak ΔF/F) HNK vs Naloxone + HNK | n = 20, 10 | Unpaired t test | $P < 0.0001$ |
| S5k | Time to peak (s) Astrocyte vs Neuron | n = 10, 10 | Unpaired t test | $P = 0.0101$ |
| S5l | AUC(a.u.) Astrocyte vs Neuron | n = 10, 10 | Unpaired t test | $P = 0.0163$ |
| S6b | (FST) Ctrl vs GFAP-ChR2 | n = 12, 11 | Ordinary one-way ANOVA | $P < 0.0001$ |
| | (FST) Ctrl vs GFAP-NpHR | n = 12, 7 | Ordinary one-way ANOVA | $P = 0.0029$ |
| | (FST) HNK vs GFAP-ChR2+HNK | n = 5, 6 | Ordinary one-way ANOVA | $P = 0.9959$ |
| | (FST) HNK vs GFAP-NpHR+HNK | n = 5, 8 | Ordinary one-way ANOVA | $P = 0.0356$ |
| | (SPT) Ctrl vs GFAP-ChR2 | n = 7, 12 | Ordinary one-way ANOVA | $P = 0.0127$ |
| | (SPT) Ctrl vs GFAP-NpHR | n = 7, 7 | Ordinary one-way ANOVA | $P = 0.0083$ |
| | (SPT) HNK vs GFAP-ChR2+HNK | n = 5, 7 | Ordinary one-way ANOVA | $P = 0.8195$ |
| | (SPT) HNK vs GFAP-NpHR+HNK | n = 5, 7 | Ordinary one-way ANOVA | $P = 0.0052$ |
| S7b | sEPSCs peak amplitude（% baseline） Ctrl vs HNK | n = 7 slices (7 cells, 2 mice) | Paired t test | $P = 0.0638$ |
| S7c | sEPSCs frequency（% baseline） Ctrl vs HNK | n = 7 slices (7 cells, 2 mice) | Paired t test | $P = 0.0556$ |
| S7f | sEPSCs peak amplitude（% baseline） GFAP-ChR2 light off vs light on | n = 20 slices (20 cells, 9 mice) | RM one-way ANOVA | $P = 0.4135$ |
| | sEPSCs peak amplitude（%baseline） GFAP-ChR2 light on vs light on+HNK | n = 20 slices (20 cells, 9 mice) | RM one-way ANOVA | $P = 0.4222$ |
| S7g | sEPSCs frequency (%baseline) GFAP-ChR2 light off vs light on | n = 20 slices (20 cells, 9 mice) | RM one-way ANOVA | $P = 0.0002$ |
| | sEPSCs frequency (%baseline) GFAP-ChR2 light on vs light | n = 20 slices (20 cells, 9 | RM one-way ANOVA | $P = 0.1886$ |

| | | | | |
|---|---|---|---|---|
| | on+HNK | mice) | | |
| S7i | sEPSCs peak amplitude (%baseline) GFAP-NpHR light off vs light on | n = 12 slices (12 cells, 6 mice) | RM one-way ANOVA | *P* = 0.9656 |
| | sEPSCs peak amplitude (%baseline) GFAP-NpHR light on vs light on+HNK | n = 12 slices (12 cells, 6 mice) | RM one-way ANOVA | *P* = 0.4796 |
| S7j | sEPSCs frequency（%baseline） GFAP-NpHR light off vs light on | n = 12 slices (12 cells, 6 mice) | RM one-way ANOVA | *P* = 0.0001 |
| | sEPSCs frequency（%baseline） GFAP-NpHR light on vs light on+HNK | n = 12 slices (12 cells, 6 mice) | RM one-way ANOVA | *P* = 0.5896 |
| S10a | (HA/Flag) Ctrl vs HNK | n = 5, 5 | Unpaired t test | *P* = 0.0265 |
| | (β-arrestin/Flag) Ctrl vs HNK | n = 5, 5 | Unpaired t test | *P* = 0.0162 |
| S10b | (HA/Flag) Ctrl vs HNK | n = 5, 5 | Unpaired t test | *P* = 0.0332 |
| | (β-arrestin/Flag) Ctrl vs HNK | n = 5, 5 | Unpaired t test | *P* = 0.0056 |
| S10c | (HA/Flag) Ctrl vs HNK | n = 5, 5 | Unpaired t test | *P* = 0.0132 |
| | (β-arrestin/Flag) Ctrl vs HNK | n = 5, 5 | Unpaired t test | *P* = 0.0017 |
| S10d | (HA/Flag) Ctrl vs HNK | n = 5, 5 | Unpaired t test | *P* = 0.0364 |
| | (β-arrestin/Flag) Ctrl vs HNK | n = 5, 5 | Unpaired t test | *P* = 0.0149 |
| S10e | (HA/Flag) Ctrl vs HNK | n = 5, 5 | Unpaired t test | *P* = 0.0044 |
| | (β-arrestin/Flag) Ctrl vs HNK | n = 5, 5 | Unpaired t test | *P* = 0.0186 |

**Table S2.**

| Protein | | Ligand | Lipid | | Water | Ions | | Box ($nm^3$) |
|---|---|---|---|---|---|---|---|---|
| μ-opioid | δ-opioid | ketamine | POPC | POPG | | $Na^+$ | $CL^-$ | |
| 1 | 1 | 1 | 240 | 60 | 39858 | 240 | 60 | 10.3*10.3*15.6 |

**Table S3.**

| REAGENT or RESOURCE | SOURCE | IDENTIFIER |
|---|---|---|
| **Animals** | | |
| C57BL/6J mice | Beijing Vital River Laboratory Animal Technology Co., Ltd. | N/A |
| GFAP-Cre | GemPharmatech Co., Ltd | T004857 |
| PV-Cre | The Jackson Laboratory | JAX.017320 |
| *Oprd1*$^{flox/flox}$ | Shanghai Model Organisms Center, Inc. | NM-CKO-220401 |
| *Oprm1*$^{flox/flox}$ | The Jackson Laboratory | JAX.030074 |
| *Oprm1*-KO | The Jackson Laboratory | JAX.007559 |
| **Chemicals, Peptides** | | |
| 2R,6R-Hydroxynorketamine hydrochloride | Tocris | 6094 |
| ketamine hydrochloride | Henry Schein, Inc. and donation from Dr. Qiang Zhou | NDC: 0404-9881 |
| MK-801 | MCE | HY-15084 |
| DAMGO | Tocris | 1171/1 |
| CTOP | Tocris | 1578 |
| DPDPE | MCE | HY-P1334 |
| ICI174864 | MCE | HY-101230 |
| Corticosterone | Biosharp | BS965 |
| CYM51010 | MCE | HY-104006 |
| MOR-DOR heterodimer allosteric inhibitor peptide | Qyaobio | 4010064769 |
| DMEM | Thermo | 12491023 |
| Neuronal Cell Medium | Gibco | 21103049 |
| Hank's Balanced Salt Solution (HBSS) | Gibco | 14175095 |
| Fetal Bovine Serum (FBS) | MeilunBio | PWL217-4 |
| Penicillin-Streptomycin | NCM | C100C5 |
| 0.25% Trypsin | NCM | C100C1 |
| Poly-D-Lysine | Sigma | P6407 |
| B27 Supplement | Gibco | A3582801 |
| GlutaMAX Supplement | Gibco | 35050061 |
| AraC | MCE | HY-13605 |
| CultureOne™ Supplement | Thermo | A3320201 |
| Phanta Max Master Mix | Vazyme | P515 |
| DMT Enzyme | TransGen Biotech | GD111-01 |
| Cell Lysis Buffer | Beyotime | P0013J |
| Lipofectamine 3000 | Thermo | L3000015 |
| CGP52432 | Millipore | SML0593 |
| Picrotoxin | MCE | HY-101391 |
| QX-314 | Tocris | 1014/100 |

| | | |
|---|---|---|
| Naloxone | Aladdin | C2304794 |
| **Critical Commercial Assays** | | |
| DNA Cleanup and Concentration Kit | Vazyme | DC301 |
| EndoFree Plasmid Maxi Kit | Vazyme | DC202 |
| BCA Protein Assay Kit | Thermo | RG235625 |
| enhanced chemiluminescence (ECL) reagent | Thermo | RJ240732 |
| **Antibodies** | | |
| Rabbit anti-GluA1 | Cell signaling technology | 13185T |
| Rabbit anti-β-Actin | Cell signaling technology | 4970T |
| mouse anti-MOR-DOR heterodimer | Kerafast, USA | EMS007 |
| rabbit anti-GFAP | Proteintech | 16825-1-AP |
| rabbit anti-MOR | Proteintech | 27625-1-AP |
| rabbit anti-DOR | Abcam | ab176324 |
| rabbit anti-GAPDH | Cell signaling technology | 5174S |
| rabbit anti-BDNF | Cell signaling technology | 47808 |
| HRP-conjugated goat anti-mouse immunoglobulin IgG | Abcam | 6789 |
| HRP-conjugated goat anti-rabbit immunoglobulin IgG | Abcam | 6721 |
| Mouse Anti-tuj1 | Biolegend | 801202 |
| Alexa Fluor 555 goat anti-mouse | Invitrogen | A21422 |
| Alexa Fluor 488 goat anti mouse | Invitrogen | 15H9L93 |
| Alexa Fluor 555 goat anti rabbit | Invitrogen | A21428 |
| ProLong$^{TM}$ Gold Antifade Mountant | Invitrogen | P36934 |
| Isotype-Matched IgG | Beyotime | A7028 |
| anti-DYKDDDDK tag recombinant antibody | Proteintech | 80801-2-RR |
| Protein A/G magnetic beads | MCE | HY-K0202 |
| anti-HA tag | Abmart | M20003S |
| anti-Flag tag | Abmart | M20008S |
| Rabbit anti-β-Arrestin 2 | Proteintech | 10171-1-AP |
| anti-GFP | Massive Photonics | N0305-DBCO |
| anti-ALFA | Massive Photonics | N0310-DBCO |
| **Virus strains** | | |
| AAV9-EF1a-DIO-EYFP | Vigenebio Bioscience(Jinan) | AV204094-AV9 |
| AAV9-EF1a-DIO-hChR2(H134R)-EYFP | Vigenebio Bioscience(Jinan) | AV201007-AV9 |
| AAV9-EF1a-DIO-eNpHR3.0-EYFP | Vigenebio Bioscience(Jinan) | AV201021-AV9 |

| | | |
|---|---|---|
| pAAV-CMV-EGFP | Made by own | N/A |
| pAAV-CMV-*Oprm1*-D147A Y148A | Made by own | N/A |
| pAAV-CMV-*Oprm*1-D147A Y148A V300A | Made by own | N/A |
| **Softwares** | | |
| Any-maze software | Stoelting | https://stoeltingco.com/Neuroscience/ANY-maze |
| ImageQuant™ TL （IQTL） ver.8 | Cytiva | https://www.cytivalifesciences.co.jp/ |
| pCLAMP 11 Standard Electrophysioloc software | Axon | PCLAMP 11 |
| FV31S-SW | OLYMPUS | lifescience.evidentscientific.com.cn/zh/downloads/detailiframe/?0[downloads][id]=847252002 |
| ImageJ | National Institutes of Health | https://imagej.nih.gov/ij/inde x.html |
| Prism | GraphPad Software | https://www.graphpad.com/ scientific-software/prism |

**Movie S1.**

HNK, but not ketamine, rapidly elevates YFP fluorescence intensity.

**Movie S2.**

MOR-DOR heterodimer maintains stable transmembrane conformation during 400 ns all-atom molecular dynamics simulation with low RMSD fluctuations.
